# A spatially resolved human brain proteome atlas for decoding brain function and disease

**DOI:** 10.1101/2025.10.16.682772

**Authors:** Qi Xiao, Yuting Xie, Meng Luo, Hui Yang, Zhengyi Yang, Ming Chen, Nan Wang, Zhaoxing Li, Jinlong Gao, Jing Yu, Wenhao Jiang, Weigang Ge, Weirong Xiang, Jianfei Lu, Zongxiang Nie, Yaoting Sun, Xinrui Lin, Wei Liu, Xu Zheng, Ning Fan, Jingyu Gu, Xu Ruyi, Fei Xu, Yunlong Zhang, Xiang-Dong Fu, Bo Li, Ed S. Lein, Tianzi Jiang, Ying Mao, Yan Li, Tiannan Guo

**Affiliations:** Affiliated Hangzhou First People’s Hospital, State Key Laboratory of Medical Proteomics, School of Medicine, School of Future Biomedicine, Westlake University, Hangzhou, Zhejiang Province, China; Westlake Center for Intelligent Proteomics, Westlake Laboratory of Life Sciences and Biomedicine, Hangzhou, Zhejiang Province, China; Research Center for Industries of the Future, School of Life Sciences, Westlake University, Hangzhou, Zhejiang Province, China; Songjiang Research Institute, Department of Emergency and Critical Disease, Songjiang Hospital Affiliated to Shanghai Jiao Tong University School of Medicine, Department-of Anatomy and Physiology, College of Basic Medical Science, Shanghai Jiao Tong University School of-Medicine, Shanghai-201600, China; Department of Neurosurgery, National Center for Neurological Disorders, Shanghai Key Laboratory of Brain Function Restoration and Neural Regeneration, Huashan Hospital, Fudan University, Shanghai 200040, China; State Key Laboratory of Medical Neurobiology and MOE MRIFrontiers Center for Brain Science, Institute for Translational Brain Research, Fudan University, Shanghai 200040, China; Beijing Key Laboratory of Brainnetome and Brain-Computer Interface, Institute of Automation, Chinese Academy of Sciences, Beijing 100190, China; Department of Anatomy, Dalian Medical University, Dalian 116044, China; Shanghai Jiao Tong University School of Medicine, 227 South Chongqing Road, Huangpu District, Shanghai 200025, China; College of Basic Medical Science South China University of Technology, Guangzhou 510641, China; Westlake Laboratory of Life Sciences and Biomedicine, Hangzhou, China; Westlake Omics Biotechnology Co., Ltd, Hangzhou, China; Xiaoxiang Institute for Brain Health and Yongzhou Central Hospital, Yongzhou 425000, China; Allen Institute for Brain Science, Seattle, 98103, Washington, USA

## Abstract

While the brain performs specialized functions across distinct regions, the spatial organization of the human brain proteome remains largely uncharted. Here we present the most comprehensive spatially-resolved human brain proteome atlas to date, comprising over 2000 post-sampling MRI registered brain sites from cortex, subcortical nuclei and cerebellum across four individuals. Our proteomic profile reveals unprecedented protein expression features in the adult human brains, such as markedly complex protein diversities compared to the transcript in the cortical regions, distinct left-right hemispheric expression differences, and sexual dimorphism in cerebellar energy metabolism. Unsupervised molecular clustering defines distinct brain territories that transcend anatomical boundaries, reflecting metabolic demands and functional specialization patterns. In addition, we collected 90 surgically resected brain tissue specimens from 49 epilepsy patients, covering 6 disease-affected regions. By integrating protein profiles from these epilepsy brain samples with our reference atlas, we identified disease-relevant protein dysregulation, like impaired astrocyte metabolism and disrupted protein homeostasis. The analyses further uncovered potential therapeutic targets including the seizure-associated purinergic receptor P2RX7. This resource depicts a high-resolution spatial proteomic landscape of the human brain, providing a foundational molecular framework for understanding brain function and disease, also the discovery of therapeutic targets.

## Introduction

The human brain enables complex cognition and behavior through intricate neural networks in distinct cortical areas and subcortical structures^1,2^. Proteins underpin these functions at every scale. Mapping the regional distribution of these proteins is essential to understanding how specific brain territories drive overall function.

To decode this molecular architecture, recent transcriptomic atlases have provided invaluable insights^3,4^, particularly in defining cell-type diversity and lineage relationships across the cortex^5,6^. While gene expression outlines the cellular blueprint, the functional state of neural tissue is further shaped by translational efficiency, protein stability, and post-translational modifications^7^. Consequently, proteomic analysis serves as a crucial complementary layer to transcriptomics. Characterizing the protein landscape, facilitates capture of the downstream functional readouts that may not be visible at the transcription level alone, offering a more holistic view of cortical specialization^8,9^.

In parallel with these molecular advances, neuroimaging has transformed our understanding of brain organization^1,10^. The Brainnetome Atlas subdivides the brain based on functional connectivity profiles^2^. This atlas has become a cornerstone for mapping the brain’s functional topography, widely used to investigate network dynamics in healthy cognition and circuit disruptions in neuropsychiatric disorders^11–15^. However, distinct functional connectivity implies distinct molecular demands. By aligning proteomic data with the Brainnetome framework, we expose unique proteomic patterns of functional brain regions. This protein-level heterogeneity diverges from the relatively uniform transcriptomic landscape of the cortex.

We present a most comprehensive spatially-resolved human brain proteome atlas to date, to bridge the gap between macroscale function and microscale composition. To improve spatial fidelity, we utilized a “post-sampling MRI registration” pipeline to ensure superior spatial precision. Unlike traditional methods that rely on approximate alignment of tissue slices, we subjected tissue punches to MRI scanning immediately after retrieval to pinpoint their exact spatial coordinates. This pipeline enables us to analyze over 2000 samples across four individuals, achieving a spatial granularity unmatched by previous studies, specifically, a 10-fold increase in cortical parcellation and a 3-fold expansion in subcortical nuclei coverage relative to existing proteome maps^4,8,9,16–18^. Crucially, our mapping of subcortical nucleus is strictly guided by anatomical atlases to ensure rigorous definition. By integrating this high-precision spatial data with PCT-DIA mass spectrometry^19^, we reveal that protein profiles strongly covary with functional connectivity patterns. This resource enables us to dissect the molecular substrate of functional networks and highlights its potential clinical utility by identifying metabolic disruptions and therapeutic targets in epilepsy.

## Results

### Brain sample collection, post-sampling MRI registration and proteomic profiling

We collected brain tissue specimens from two Chinese males and two Chinese females, with no history of neuropsychiatric or neuropathological conditions (**Fig. 1a**, **Supplementary Fig. S1a** and **Supplementary methods**). Each human brain was perfusion-fixed in formalin within ten hours postmortem. For proteomic analysis, 307–379 discrete cortical and subcortical sampling points were collected using a punch biopsy tool from both hemispheres of each brain, yielding a total of 1384 spatially resolved sample locations across the four individuals (**Supplementary Table 1**). Additionally, MRI was performed on each brain (see **Supplementary methods**) to obtain standardized spatial coordinates for sample localization (**Supplementary Table 1**). These native coordinates were then transformed to the Montreal Neurological Institute (MNI) coordinate space^20^, enabling cross-brain comparisons and integration with public databases. Samples from the cortex was assigned to 186 subregions defined by the Brainnetome Atlas^2^, which provides detailed parcellation based on both structural connectivity and functional activation patterns. This MRI post-sampling registration approach represents a critical advance over previous brain proteomics studies ^4,8,16^, enabling direct correlation between molecular signatures and functionally defined brain territories (**Supplementary Fig. S1b**).

**Figure 1.**
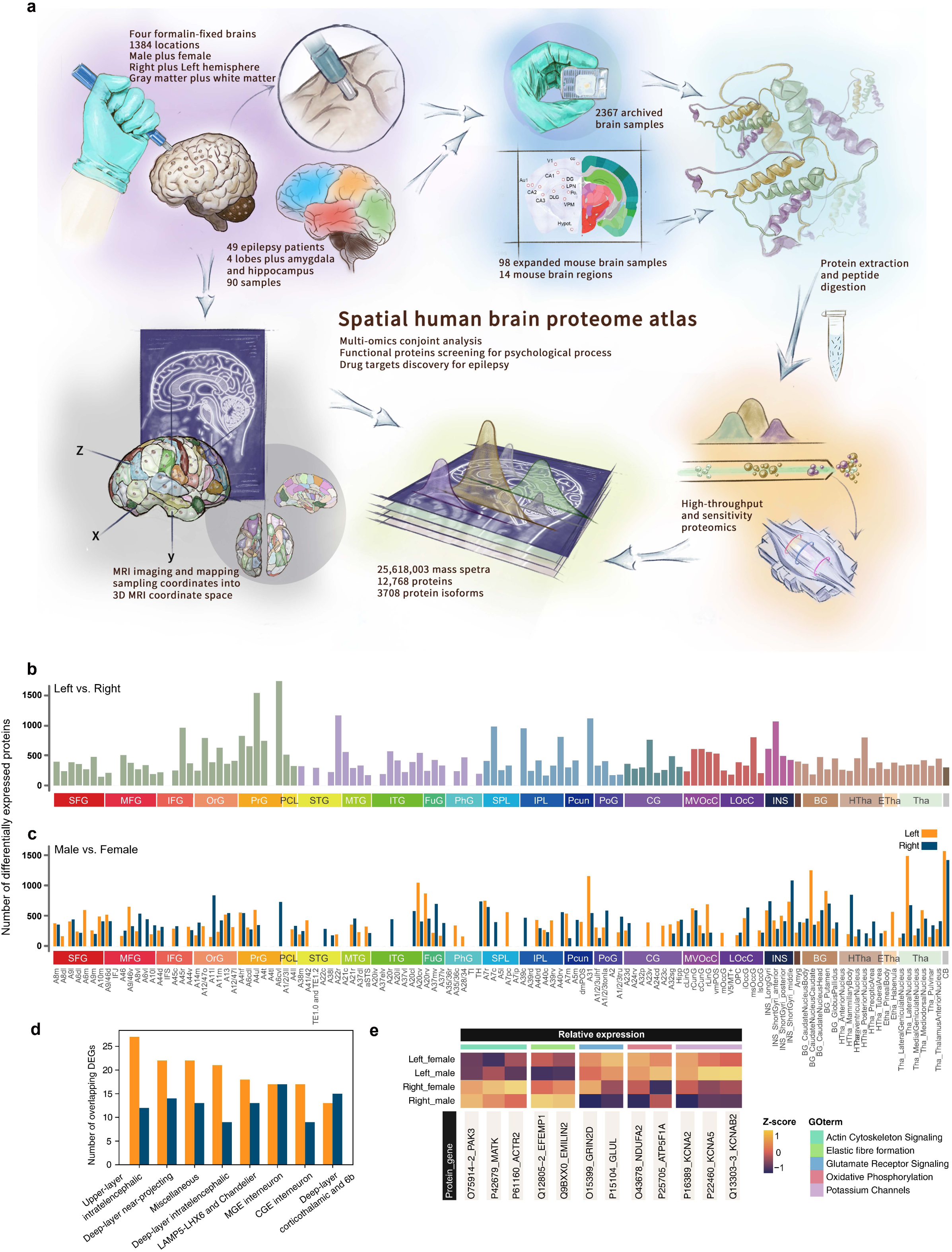
Generation and characterization of the human brain proteome atlas. **a,** Flowchart illustrating the comprehensive pipeline for generating the draft human brain proteome atlas. **b,** Bar plot depicting the number of significantly dysregulated proteins between left and right hemispheres (Student’s t-test, Benjamini-Hochberg adjusted p < 0.05). Colors represent different brain regions: frontal lobe (FL), temporal lobe (TL), parietal lobe (PL), limbic lobe (LL), occipital lobe (OL), insular lobe (IL), subcortical nuclei, and cerebellum. **c,** Bar plot showing the number of significantly dysregulated proteins of male vs. female subjects (Student’s t-test, Benjamini-Hochberg adjusted p < 0.05). Orange bars represent left hemisphere regions; dark blue bars represent right hemisphere regions. **d,** The number of Left (orange) and Right (dark blue) overlapped DEGs is shown for each neuronal cell type, ordered by the number of Left M1 DEGs. **e,** Heatmap displaying protein expression patterns for significantly dysregulated proteins in the PrG. Color scale represents z-score normalized protein expression levels, with orange indicating higher expression and dark purple indicating lower expression relative to the mean.

To enhance spatial coverage of deep brain structures, we expanded sampling to include additional subcortical nuclei and the cerebellum. These regions were anatomically annotated according to the Paxinos and Watson human brain atlas^21^. The cortical regions were further separated into grey and white matter compartments, and selected regions was sampled in biological replicates, resulting in 2277 spatially resolved samples in total. In addition, we collected 90 brain tissue specimens from 49 individuals with epilepsy, spanning six regions implicated in seizure disorders: the frontal, temporal, parietal, and occipital lobes, as well as the amygdala and hippocampus (**Supplementary Table 1**).

We employed PCT coupled with DIA-MS for proteomic analysis of human brain tissue^19,22,23^. First, we built a spectral library that is the most comprehensive to date for human brain, containing 196,892 peptides from 12,768 proteins, which includes 3708 protein isoforms (**Supplementary Fig.2 a-h**). Subsequently, all samples were analyzed using a high-resolution MS, which quantified 171,225 unique peptides corresponding to 11,772 protein groups and 3295 protein isoforms. This comprehensive detection of protein isoforms across brain regions revealed extensive alternative splicing diversity. Plectin (PLEC) exemplified striking region-specific isoform expression, with PLEC 1e enriched in the cerebral cortex (1.2-fold) and PLEC1 enriched in subcortical nuclei (1.9-fold), reflecting different cytoskeletal requirements between these anatomically distinct brain regions^24^ (**Supplementary Fig.4a and b)**.

To monitor and ensure data quality, we implemented multiple quality control checkpoints throughout the brain proteomics pipeline (**Supplementary Fig.1c** and **Supplementary Methods**). Median coefficient of variation was 1% for technical replicates and 2.6% for biological replicates from matched brain regions (**Supplementary Fig.2i**). The dynamic range spanned seven orders of magnitude, from histone protein to mRNA splicing molecules. Principal component analysis revealed that biological variables (brain region, sex, individual) explained 32.2% of total variance, while technical factors contributed 1.1% (**Supplementary Fig.1e and f**).

### Protein expression signatures underlying cortical hemispheric specialization

Analysis of hemispheric protein expression patterns provides molecular insights into functional brain lateralization. A paired comparison of 121 left and right hemisphere samples revealed widespread hemispheric differences in protein abundance. Notably, the precentral gyrus (PrG; primary motor cortex, M1 in Brodmann area 4) displayed the most pronounced asymmetry (**Fig. 1b**), with 1725 differentially expressed proteins (DEPs) identified in the caudal ventrolateral region (p < 0.05).

The protein profile of the left M1 appears to be characterized by an enrichment of components involved in modulating firing kinetics and meeting bioenergetic demands. This profile includes voltage-gated potassium channel subunits (KCNA2, KCNA5, KCNAB2), mitochondrial respiratory chain components (NDUFA2, ATP5F1A), and key metabolic enzymes (PDHB, GLUL) (**Fig. 1e**). In contrast, the right hemisphere showed preferential expression of components associated with structural plasticity and extracellular matrix integrity (PAK3, EFEMP1, EMILIN2), collectively facilitating extracellular microenvironment organization and providing the structural scaffolding essential for synaptic stabilization.

While the upregulation of potassium channels might typically be associated with hyperpolarization and reduced excitability, the specific localization of KCNA (Kv1 family) channels to the axon initial segment (AIS) suggests a more nuanced role in enhancing spike timing precision and filtering synaptic noise^25^. Rather than pointing to a simple increase in baseline excitability, these data suggest an optimized molecular infrastructure designed for high-frequency, temporally faithful signaling. This interpretation is further supported by the presence of synaptic vesicle proteins such as SYN1 and SNAP91 in our broader dataset, which implies a capacity for reliable neurotransmitter release during the complex motor sequences managed by the dominant hemisphere.

To explore the potential cellular substrates of this asymmetry, we performed an overlap analysis between the proteomic DEPs and transcriptomic cell-type signatures from the human brain atlas^5^. By intersecting these feature sets, we inferred potential associations between specific cell lineages and the observed hemispheric protein landscapes.

The Left M1 proteome showed a notable overlap with signatures of superficial-layer intratelencephalic (IT) neurons, hinting that the dominant hemisphere may prioritize intracortical communication and sensorimotor integration (**Fig. 1d**). The enrichment of KCNA channels in the left hemisphere is notable in this context. These channels are prominently localized to the AIS of pyramidal neurons^25^, where they contribute to filtering synaptic noise and enhancing spike timing precision^26^. Their upregulation is therefore consistent with a model in which the dominant hemisphere requires greater temporal fidelity of neuronal firing. Furthermore, the co-enrichment of mitochondrial proteins (NDUFA2) suggests that these demands are met with a corresponding bioenergetic support^27^.

These hemispheric differences in protein expression provide molecular evidence for the functional specialization of motor cortical regions and suggest that lateralized motor behaviours may be supported by distinct underlying proteome landscapes that optimize each hemisphere for its specialized computational demands.

### Post-transcriptional regulation drives regional proteomic heterogeneity in the cerebral cortex

To systematically characterize molecular diversity across the human brain, we integrated proteomic and transcriptomic profiles mapped to the 121 regions of the Brainnetome Atlas^2^. Left and right hemisphere proteomic data were consolidated to ensure spatial alignment with available transcriptomic datasets. By employing a protein enrichment scoring algorithm^28,29^, we quantified regional differentially expressed (RDE) proteins and transcripts to facilitate direct multi-omics comparison. This approach identified an average of 428 RDE proteins per region **(Supplementary Fig. S3)**. To evaluate the concordance between the proteome and transcriptome, we applied the same specificity metric to spatially matched public transcriptomic data^3^. We observed a striking divergence in the cerebral cortex: the number of RDE proteins significantly exceeded that of RDE transcripts, with the average count of cortical RDE proteins being approximately 1.8-fold (320 *vs* 175) higher than that of transcripts **(Fig. 2a, b)**. While previous studies have emphasized the relative transcriptional homogeneity of the cortex^3^, our data reveal that cortical heterogeneity is markedly amplified at the protein level. This suggests that the distinct functional demands of specialized cortical areas are governed by region-specific post-transcriptional regulation rather than transcriptional output alone.

**Figure 2.**
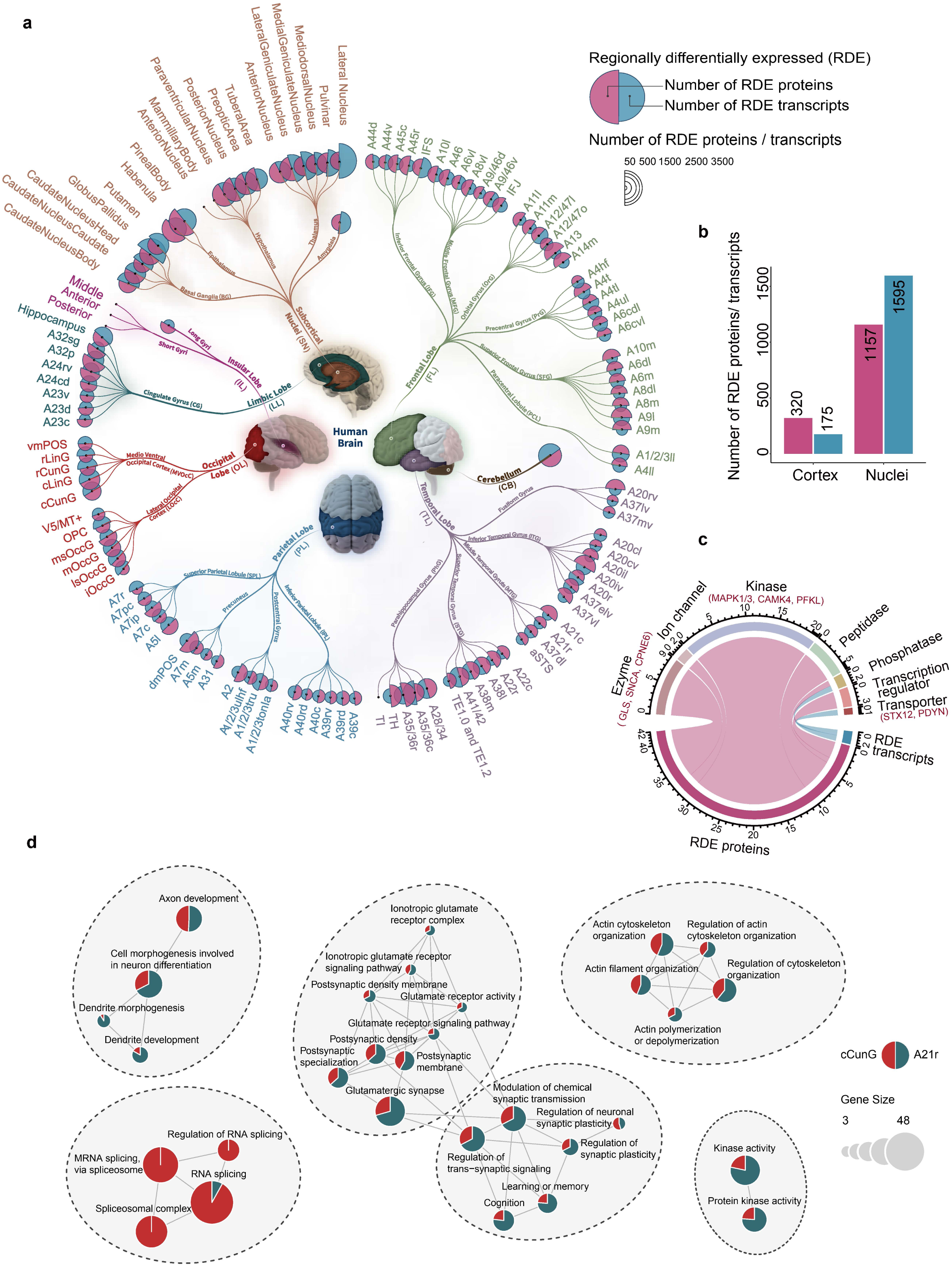
Regional expression patterns and functional characterization of proteins and transcripts across brain subregions. **a,** Comparative analysis of regionally differentially expressed (RDE) proteins and transcripts across 121 brain subregions. Each pie chart represents a single subregion; chart size corresponds to the total number of RDE proteins (specificity score > 0.93, 95th percentile threshold; ANOVA with Benjamini-Hochberg adjustment, $p < 0.05$) and RDE transcripts (specificity score > 0.94, 95th percentile threshold; ANOVA with Benjamini-Hochberg adjustment, $p < 0.05$) identified in that region. **b,** Quantification of RDE proteins (pink) and RDE transcripts (blue) in the cerebral cortex versus subcortical nuclei. Values above bars denote the mean counts across all regions within each category. **c,** Chord diagram illustrating the distribution of drug targets among RDE proteins and transcripts specific to the A21r region. Drug target data were retrieved from the ChEMBL database (accessed April 25, 2025). **d,** Gene Ontology (GO) pathway enrichment analysis for cCunG (visual association area) and A21r (middle temporal gyrus) RDE proteins and transcripts. Red bars denote pathways specifically enriched in RDE proteins; green bars represent pathways specifically enriched in RDE transcripts.

We specifically focused on the rostral Middle Temporal Gyrus (A21r), identifying it as a region of dual clinical and functional significance. Clinically, A21r is a frequent neocortical focus in temporal lobe epilepsy; functionally, it serves as a critical node within the Default Mode Network, playing a pivotal role in semantic processing^30,31^. By comparing A21r against 120 other cortical regions, we defined a distinct molecular signature comprising approximately 104 RDE proteins and 10 RDE transcripts. Notably, we observed a marked discrepancy between the transcriptome and proteome. For example, while specific transcripts, such as PDYN and NPTXR, were identified as RDEs due to their high abundance, their corresponding protein levels were not proportionally elevated. This divergence suggests that post-transcriptional or post-translational regulatory mechanisms heavily influence the final proteomic landscape of A21r.

Functional annotation of the proteins preferentially expressed in A21r indicated a molecular profile characterized by kinase activities, which represented the most enriched functional category (**Fig. 2c**). This signature included proteins associated with signal transduction and synaptic plasticity, such as MAPK1/3 and CAMK4, as well as the calcium-dependent membrane-binding protein CPNE6. Concurrently, we observed an enrichment of metabolic enzymes, specifically the rate-limiting glycolytic enzyme PFKL and the glutamine-hydrolyzing enzyme GLS, alongside proteins mediating vesicle trafficking such as SNCA and STX12. These data associate the A21r with selectively elevated signaling capacity and metabolic demand, physiological features that are potentially relevant to the region’s intrinsic susceptibility to hyperexcitability.

To contextualize the molecular features of A21r, we performed a comparative pathway enrichment analysis against one reference regions selected for their distinct functional profiles (**Fig. 2d**), the Primary Visual Cortex (V1, represented by cCunG). V1 was selected as a baseline representing early sensory cortex, characterized by distinct laminar architecture and functional specialization for sensory processing^33^. Notably, fundamental structural pathways, such as actin cytoskeleton organization, showed comparable enrichment levels across both A21r and V1, suggesting that their functional differences are not attributable to variations in the abundance of basic structural components, but rather to distinct regulatory programs.

A striking molecular divergence was observed between these sensory and association cortices. V1 exhibited specific enrichment in pathways related to RNA splicing and mRNA metabolism. This aligns with the biological requirement for high proteomic diversity in sensory cortex, where alternative splicing generates isoform diversity in cell adhesion molecules and ion channels to facilitate precise circuit assembly^35^. In contrast, A21r was characterized by an enrichment in pathways governing nervous system development and the regulation of neuronal synaptic plasticity via kinase activity. While V1 prioritizes splice-variant diversity for structural specification, the A21r signature indicates a mechanism reliant on kinase-mediated signalling and developmental pathways. While comparable enrichment of actin cytoskeleton components indicates a shared foundation of structural integrity, divergent pathways—RNA processing in V1 versus kinase-mediated signaling in A21r—point to distinct operational strategies. Primary sensory cortices prioritize robust circuit assembly via isoform diversity, whereas association cortices favor a proteomic architecture optimized for rapid, reversible modulation of synaptic weight.

### Sex-specific molecular signatures in the cerebellum

Given the rarity of matched male and female tissue samples, we investigated whether sex-specific molecular signatures persist across the brain. We focused our pairwise proteomic analysis on the cerebellum, which displayed marked molecular sexual dimorphism, and discovered distinct metabolic and cellular strategies between sexes (**Fig. 1c**).

Females demonstrated enhanced mitochondrial efficiency through the upregulation of fatty acid metabolism enzymes (CPT1A, FAH, HADHA, and ACADS). In contrast, males exhibited rapid energy production via enhanced glycolytic activity and mitochondrial biogenesis (NMNAT1, ATP1A4, and MRPS12/16/23). Notably, this male-specific metabolic profile was accompanied by heightened neuroinflammatory signaling, characterized by complement system activation (C4B, CFH, C1QA) and pro-inflammatory mediators (IFI44L, TNFRSF21, DAXX) (**Supplementary Fig. S2l**). These divergent profiles suggest that the cerebellum possesses sex-specific vulnerabilities to metabolic stress and neuroinflammation.

### Connective hub regions exhibit specific molecular architecture

Brain regions vary in their connectivity patterns, with some regions serving as highly connected “hubs” that integrate information across multiple brain networks. To examine how protein expression relates to brain intrinsic connectional architecture, we analyzed its specialization across regions with different connectivity profiles.

We first examined connectivity-based molecular specialization. We quantified functional connectivity (FC) degrees for each of the 184 cortical regions by summing their strongly connected neighbors, using functional connectivity matrices derived from HCP resting-state fMRI data with matched cortical parcellation^2^ (**Fig. 3a and Supplementary Fig. S5a**). To delineate the molecular architecture underlying these connectivity profiles, we integrated FC degrees with multi-level molecular datasets encompassing grey matter (GM) proteomics, white matter (WM) proteomics, and GM transcriptomics (**Fig. 3c**). Through a stringent multi-level filtering strategy (Spearman correlation, FDR < 0.2, and top 10% effect size), we delineated a unique molecular signature specific to high-connectivity brain hubs, uncovering a fundamental biophysical trade-off: highly connected regions prioritize molecular quality control and metabolic maintenance over the density of electrophysiological signaling hardware.

**Figure 3.**
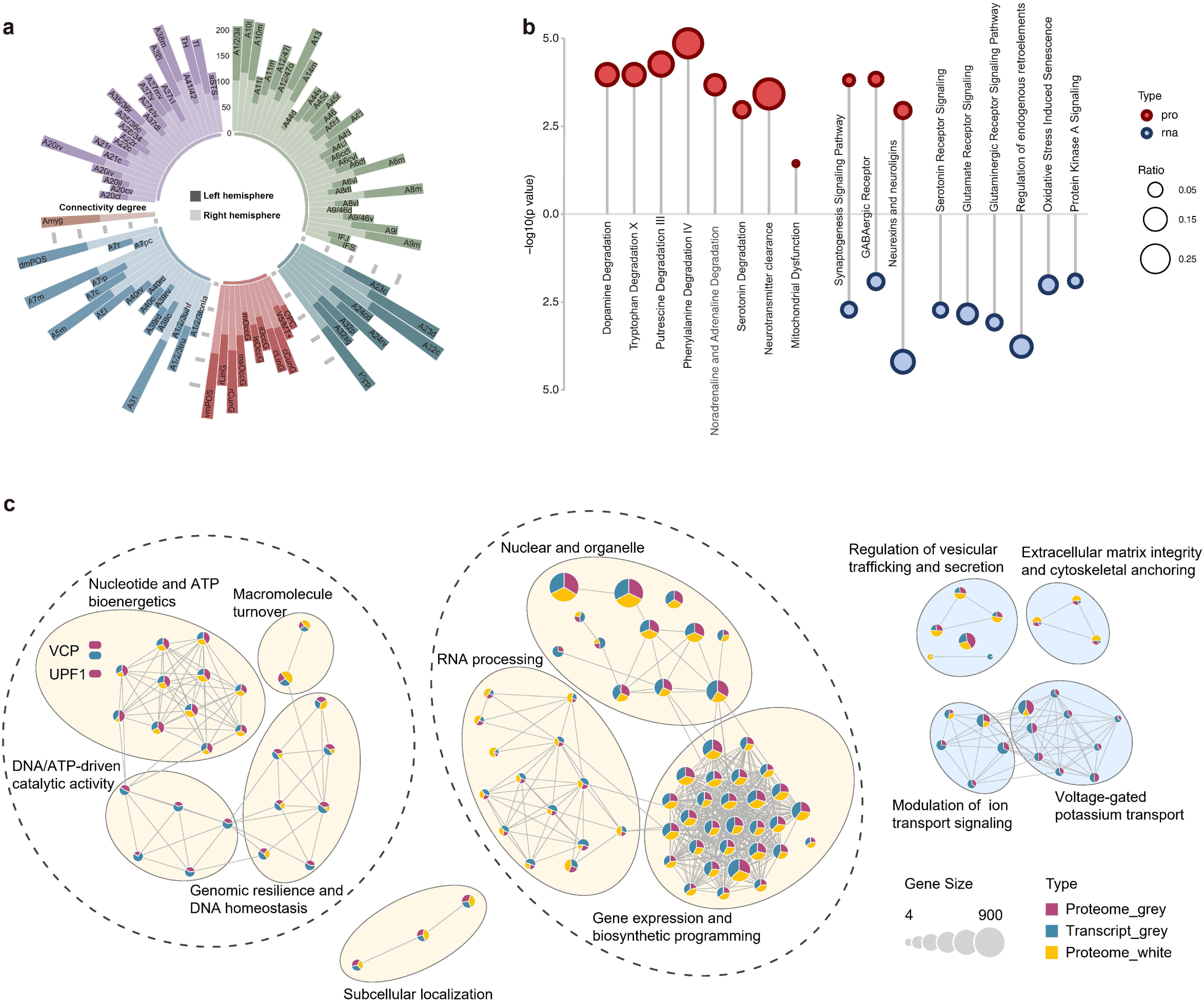
Integration of proteomics data with brain structural connectome analysis. **a,** Bar plot shows connectivity degrees for each cortical region, qua ntified by summing strongly connected neighbors using functional connectivity matrices derived from HCP resting-state fMRI data with matched cortical parcellation. Dark bars represent left hemisphere regions; light bars represent right hemisphere regions. Higher values indicate greater functional connectivity with neighboring regions. **b,** Ingenuity Pathway Analysis (IPA) of genes positively correlated with connectivity degree. Red and blue circles denote pathways enriched in positively correlated proteins and transcripts, respectively. **c,** Pathway enrichment analysis comparing grey and white matter. Red and blue bars represent pathways enriched in proteins and transcripts positively correlated with connectivity in grey matter, respectively; yellow bars represent pathways specifically enriched in proteins positively correlated in white matter.

This highly connected “hubs” is supported by a positive correlation network hierarchically organized into functional modules of bioenergetic and surveillance machinery. At the core, modules dedicated to nucleotide and ATP bioenergetics provide the metabolic foundation, fueling a sophisticated system of RNA processing and macromolecule turnover characterized by hub proteins VCP and UPF1. In the grey matter, this suggests that hubs maintain connectivity not by increasing protein quantity, but by accelerating the turnover of damaged components and ensuring high-fidelity translation through the nonsense-mediated decay pathway. Notably, the prominence of RNA-binding modules in the white matter proteomics suggests that long-distance connectivity is further supported by robust local translation within axons to maintain structural integrity far from the soma. In contrast, proteins associated with electrophysiological excitability and structural scaffolding—such as inhibitory potassium channels (KCNC3) and basic signaling hardware (SCN1A, ATP2B1/2)—exhibited negative correlations with connectivity. This indicates that hubs adopt a “lean” signaling strategy, reducing the energetic cost of baseline ion pumping and lowering firing thresholds to facilitate rapid synchronization.

While hubs minimize their permanent structural hardware, they maintain the temporal precision required for high-order integration through the efficient clearance of synaptic signals. This necessity is reflected at the protein level by the specialized expression of neurotransmitter-degrading enzymes—including ALDH2, MAOA, and MAOB—which ensure rapid signal termination to prepare for the next wave of information (**Fig. 3b**). Crucially, we observed a distinct decoupling between high mRNA abundance and actual protein translation in key metabolic and synaptic pathways. By maintaining a vast “transcriptional priming” state— evidenced by a high mRNA-to-protein ratio—hubs preserve a ready-to-deploy molecular blueprint. This allows the system to avoid the metabolic burden of maintaining a dense protein infrastructure while retaining the capacity for rapid, on-demand synaptic remodeling and the efficient signaling flux essential for complex cognitive processing.

### Multi-omics clustering reveals 21 molecularly discrete brain territories

The creation of a comprehensive molecular atlas is essential for deciphering how the brain’s rigid anatomical scaffold supports its vast functional plasticity and susceptibility to disease. While traditional atlases categorize brain regions based on cytoarchitecture, a multi-layered molecular map provides a dynamic readout of the physiological state and therapeutic potential of these territories. To systematically identify molecular territories that transcend classical anatomical boundaries, we performed unsupervised multi-omics clustering on spatially matched proteomic and transcriptomic datasets covering brain regions (**Fig. 4a**). Following independent normalization and batch effect correction for each omics layer, we employed the MOVICS framework^36^, which aggregates consensus clustering across multiple algorithms. The optimal cluster number of k=21 was determined based on silhouette width and Clustering Prediction Index (**Supplementary Fig. S6**).

**Figure 4.**
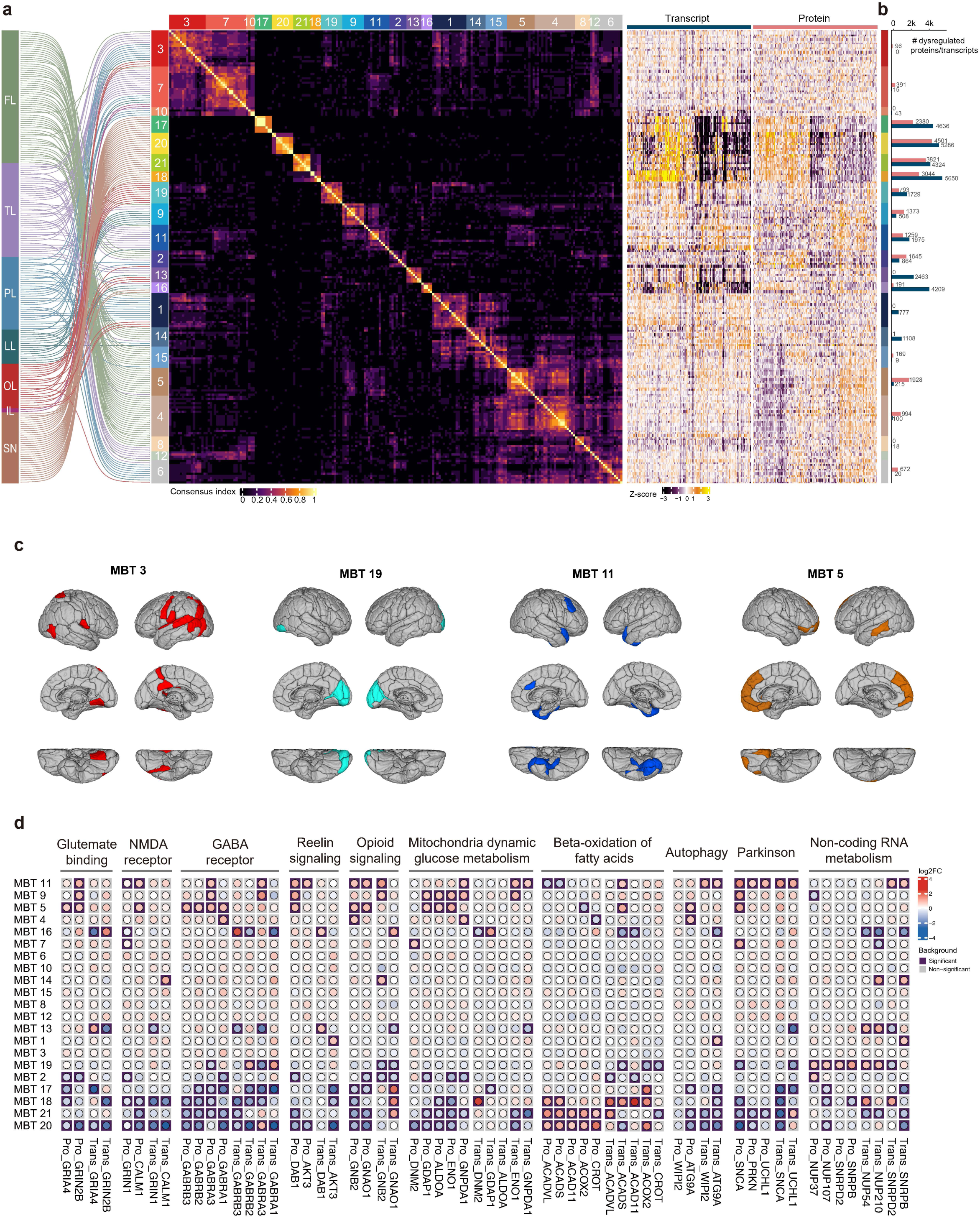
Molecular-based brain parcellation reveals functional organization patterns. **a,** Consensus clustering of 216 brain subregions into 21 MBTs. The consensus heatmap displays the consensus index for each subregion. **b,** Bar plot displaying the number of differentially expressed proteins (red) and transcripts (dark blue) for each of the 21 MBTs. Each cluster compared to all other clusters combined. Significance thresholds were set at Benjamini-Hochberg adjusted p < 0.05 and |fold-change| > 1.5. **c,** Three-dimensional visualization of MBTs mapped onto standard brain anatomy. Views shown include lateral, medial, and ventral perspectives of both hemispheres. **d,** Heatmap showing differential gene expression profiles across 21 MBTs. Each row represents an MBT and each column represents a gene. Values indicate log2 fold change of gene expression in each MBT compared to all other MBTs combined. Red indicates upregulation and blue indicates downregulation.

The 21 Molecular Brain Territories (MBTs) revealed both alignment with classical networks and novel organizational principles. For example, MBT 19 completely overlapped with the visual network, while MBTs 5 and 9 corresponded to limbic system components (**Fig. 4c**). MBT 11 encompassed regions of the central executive network, including entorhinal cortex, cingulate areas, and Brodmann areas 20 and 38—regions earliest affected in Alzheimer’s disease (AD). MBT 21 consisted exclusively of thalamic nuclei (**Supplementary table 4**). Behavioral domain analysis using BrainMap metadata revealed distinct functional specializations^37^: MBT 5 showed strong enrichment for emotion and cognition, MBT 19 for vision, and MBT 9 for cognition (**Supplementary Fig. S7)**. Cell-type enrichment analysis^38^ demonstrated that MBTs 5 and 11 were dominated by glutamatergic neurons, while MBT 19 showed mixed GABAergic and glutamatergic populations.

### Distinct molecular patterns define functional specialization across brain territories

Pathway analysis revealed how distinct molecular patterns align with regional functional specialization (**Fig. 4b and S8**). Limbic and executive territories (MBTs 5, 9, 11) supporting emotion, cognition, and executive control maintained high expression of neurotransmitter diversity and synaptic plasticity machinery. These regions showed coordinated upregulation of serotonin signaling proteins (AKT3, BDNF, BRAF)—critical for mood regulation in limbic circuits—alongside glutamatergic components (GRIA4, GRIN2A, GRIN2B) essential for cognitive flexibility (**Fig. 4d**). MBT 5, encompassing emotional processing regions, uniquely expressed high levels of GABAergic receptors (GABRA1, GABRA3, GABRB2, GABRB3), reflecting the inhibitory control required for emotional regulation. The sustained expression of reelin signaling and synaptic plasticity pathways in these territories supports their need for continuous adaptation to complex cognitive and emotional demands. This molecular complexity was matched with high metabolic investment. MBTs 5, 9 and 11 showed enhanced glucose metabolism proteins (ALDOA, ENO1, GNPDA1) and mitochondrial fission machinery (DNM2, GDAP1, MCU), reflecting the energetic cost of maintaining diverse neurotransmitter systems and plasticity mechanisms. MBT 11, which encompasses regions vulnerable to AD, also showed elevated expression of pathways associated with Parkinson’s and Huntington’s diseases. This suggests that the high neurotransmission demands and associated metabolic stress required to maintain cognitive flexibility may contribute to the heightened susceptibility of these regions to neurodegeneration. Supporting this, MBTs 9 and 11 exhibited upregulation of autophagy markers in microglia and astrocytes at the transcriptional level, reflecting unstable cellular stress responses may contribute to neurodegeneration.

In striking contrast, MBT 19 (visual network) exhibited a molecularly streamlined architecture befitting its specialized sensory function. This territory showed systematic downregulation of neurotransmitter and metabolic pathways while selectively maintaining RNA processing machinery (nucleoporins and snRNPs). This molecular profile suggests that primary sensory regions achieve functional efficiency through molecular specialization rather than flexibility, potentially explaining their relative resilience to age-related decline^39,40^.

Subcortical territories displayed distinct metabolic adaptations reflecting their functional roles. MBTs 20 and 21, encompassing thalamus, epithalamus, and hypothalamus, showed preferential expression of fatty acid oxidation and peroxisomal lipid metabolism pathways, consistent with the metabolic homeostasis and circadian regulation functions of these regions. (**Supplementary Fig. S8**).

These molecular territories reveal a fundamental principle: brain regions optimize the metabolic cost of molecular machinery to match functional needs—cognitive regions pay for flexibility, sensory regions streamline for efficiency.

### Brain proteome atlas reveals extensive molecular perturbations in epilepsy

To demonstrate the translational value of our brain proteome atlas, we analyzed 90 brain tissue samples from 49 epilepsy patients across six regions: frontal, parietal, temporal, and occipital lobes, plus amygdala and hippocampus (**Fig. 5a**). Obtaining healthy brain tissue for comparison remains challenging due to limited post-mortem availability and ethical constraints, making our reference atlas particularly valuable. Comparative analysis identified 2381 differentially expressed proteins (|Fold change| > 3, adjusted p < 0.05, **Supplementary Fig. S9a and b**), with 969 proteins consistently dysregulated across all regions. These conserved changes enriched for acute phase response, Rho family GTPases and axonal guidance pathways (**Supplementary Fig. S9c and d**), revealing core molecular perturbations in epilepsy.

**Figure 5.**
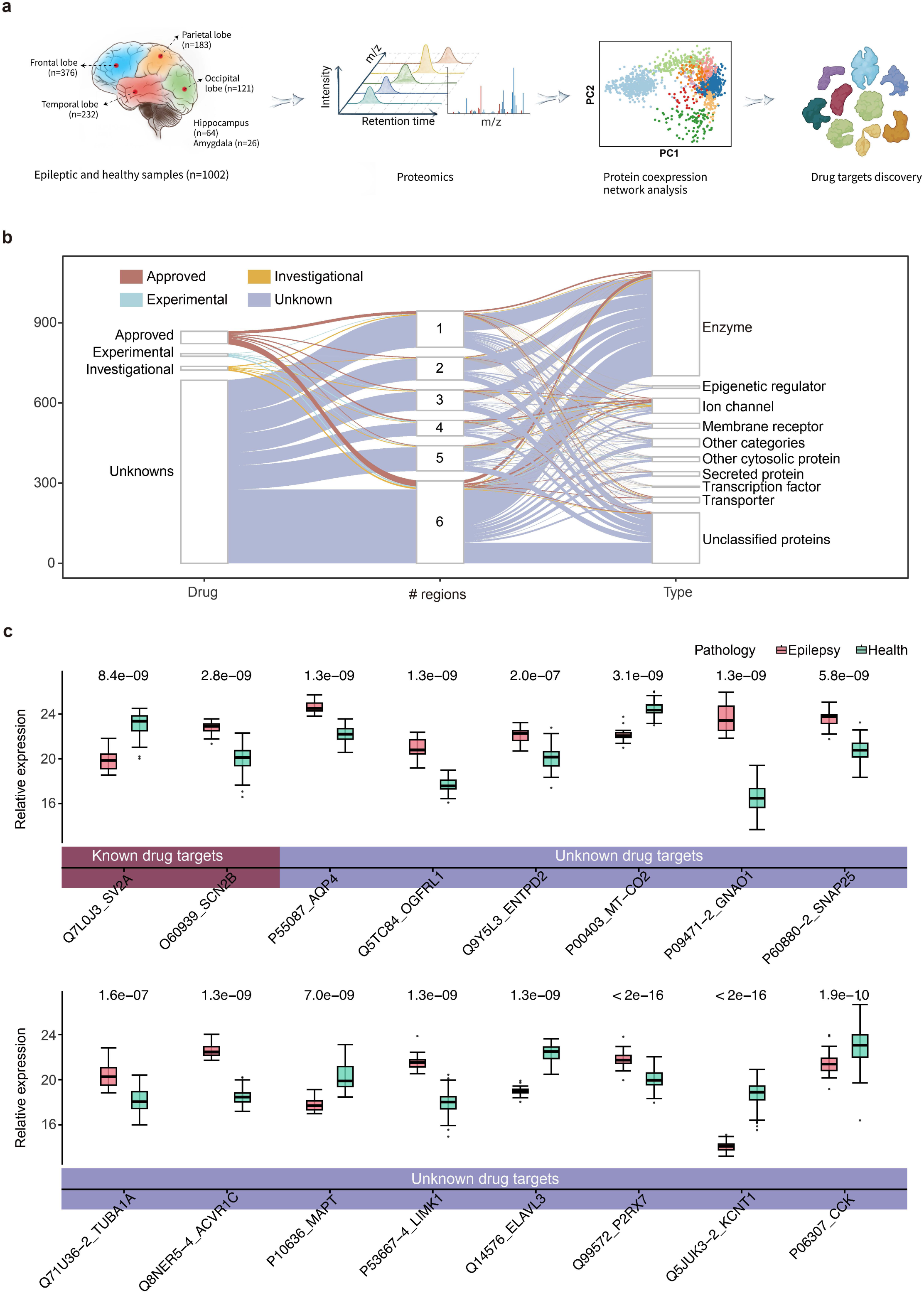
Anti-epilepsy drug target screening through region-specific dysregulation analysis. **a,** Pipeline for epilepsy drug target discovery. Epileptic brain proteome data were integrated with corresponding brain region data from our healthy brain database, yielding a total of 1002 samples for analysis. Protein expression matrices from brain tissues of healthy individuals and epileptic patients were analyzed using WGCNA, specifically comparing healthy temporal lobe versus epileptic temporal lobe samples. **b,** Sankey diagram illustrating the regional distribution and drug target classification of 736 potential drug targets, including 51 proteins that are targets of FDA-approved anti-epileptic drugs. The flow shows: (left) dysregulated potential drug targets, (middle) number of brain regions where each protein is dysregulated, and (right) drug target classification. Drug target annotations were obtained from ChEMBL and DrugBank databases (ChEMBL downloaded April 25, 2025; DrugBank downloaded April 29, 2025). **c,** Relative expression of identified drug targets in temporal lobe patients. Box plots display the relative expression levels (log₂ intensity from mass spectrometry) of 16 candidate drug targets comparing epileptic versus healthy temporal lobe and hippocampus samples. Wilcoxon test Benjamini-Hochberg adjusted p-values are labeled above each comparison. Targets with brown background represent the three proteins that are targets of currently approved anti-epileptic drugs.

To assess therapeutic potential, we cross-referenced dysregulated proteins with ChEMBL^41^ and DrugBank^42^ databases, identifying 736 potential drug targets (**Fig. 5b**). Notably, 51 of these proteins are already targeted by FDA-approved anti-epileptic drugs, validating our approach. The remaining 685 unexplored targets represent substantial opportunities for expanding the therapeutic arsenal against epilepsy.

### Brain proteome atlas facilitates prioritization of epilepsy drug targets

To identify mechanism-based targets for temporal lobe epilepsy (TLE), we performed weighted correlation network analysis (WGCNA) on TLE patient samples, revealing seven co-expression modules (**Supplementary Fig. S10a**). Comparison with healthy temporal lobe networks revealed poor preservation of two modules: Module 4 (integrin cell surface interactions) and Module 5 (fatty acid β-oxidation), both enriched for astrocyte markers (**Supplementary Fig. S10b**). This disruption of integrin signaling and metabolic homeostasis in astrocytes—key regulators of seizure propagation and neuroinflammation—suggests novel therapeutic avenues. To enhance the spatial resolution of our therapeutic discovery, we extended our WGCNA framework to include hippocampal (Hipp) tissues from TLE patients. Comparative network analysis identified seven modules in the Hipp network, four of which (M3, M4, M5, and M7) exhibited poor preservation compared to healthy controls (**Supplementary Fig. S10c**).

Based on these network disruptions, we prioritized drug targets using stringent criteria for TLE (**Supplementary Fig. S10d**): (i) significant differential expression in TL or Hipp, (ii) brain specific expression proteins, (iii) members of the disrupted modules identified in either region (TL M4/5 or Hipp M3/4/5/7), and (iv) druggability in ChEMBL/DrugBank. This approach identified 16 candidate targets, including two validated by current anti-epileptic drugs and 14 novel targets (**Fig. 5c, Supplementary Table 7**).

Among the identified candidates, SV2A (synaptic vesicle glycoprotein 2A) and SCN2B (sodium voltage-gated channel beta subunit) were prominently featured as key upregulated targets. Given that both are established, FDA-approved targets in epilepsy therapy, their successful capture highlights the clinical relevance and high precision of our spatial proteomic mapping. Beyond these established markers, our analysis also identified P2RX7 (purinergic receptor P2X7) as a notable upregulated target. The P2RX7 receptor is well-documented in its role as a driver of seizure-induced neuroinflammation, acting through the release of IL-1β, microglia activation, and the disruption of the blood-brain barrier^43,44^. Consequently, its identification within our dataset serves as a robust proof-of-principle for our analytical pipeline. Together, these findings demonstrate that integrating spatial proteomics with network analysis can move beyond symptomatic treatment to identify disease-driving molecular networks, facilitating both the validation of existing therapeutic anchors and the discovery of novel interventions, such as the repurposing of P2RX7 inhibitors (e.g., AZD-9056, CE-224535, and JNJ-5417544) for refractory epilepsy^42^.

## Discussion

The orchestration of human brain function requires a precise alignment between its macroscale connectivity and microscale molecular composition. In this study, we present the most comprehensive spatially-resolved human brain proteome atlas to date, bridging a critical gap in our understanding of the brain’s molecular architecture. By employing a high-precision MRI post-sampling registration pipeline, we achieve a spatial granularity that surpasses previous efforts. To ensure the technical fidelity and reproducibility of this large-scale dataset, we leveraged PCT-assisted proteomics, which provided the standardized and high-throughput protein extraction necessary for processing over 2,000 micro-scale tissue specimens with high consistency. Importantly, the mechanical force of pressure cycling facilitated the enhanced solubilization of hydrophobic membrane-associated proteins, enabling a deep coverage of the brain’s signalling machinery that are often underrepresented in traditional lysis protocols. This high-resolution mapping reveals that the proteomic landscape of the human cortex is far more diverse than its transcriptomic counterpart, suggesting that the functional specialization of the brain is heavily shaped by post-transcriptional regulatory programs.

A central finding of our work is that protein-level molecular diversity in the cerebral cortex is approximately 1.8-fold higher than that observed at the transcript level (**Fig. 2a**). While previous transcriptomic studies have highlighted the relative homogeneity of the human cortex compared to other organs, our data suggest that cortical areas achieve functional specialization through robust post-transcriptional and post-translational regulatory programs. The striking discrepancy observed in the rostral Middle Temporal Gyrus (A21r), where high transcript levels of markers like PDYN and NPTXR do not translate into proportional protein abundance, exemplifies this regulatory decoupling. Such mechanisms, potentially involving mRNA stability, translational efficiency, or protein turnover, allow the brain to fine-tune its signalling and metabolic machinery to meet the specific demands of association versus sensory cortex.

By integrating our proteomic map with the Brainnetome framework, we provide a molecular basis for the brain’s functional topography. We demonstrate that functional connectivity “hubs”, critical nodes for information integration, are characterized by specialized proteomic signatures, including elevated expression of neurotransmitter-degrading enzymes (e.g., ALDH2, MAOA/B) and synaptic adhesion molecules. This finding suggests the high-degree connectivity must maintain superior neurotransmitter clearance and synaptic stability to sustain intense information flow.

Our characterization of 21 MBTs reveals that molecular boundaries often transcend classical cytoarchitectural divisions, uncovering fundamental principles governing disease susceptibility. We observed a critical trade-off between functional flexibility and metabolic sustainability: territories supporting cognition and emotion maintain metabolically expensive machinery—including diverse neurotransmitter systems and plasticity pathways—which creates inherent vulnerability to neurodegeneration. This vulnerability is exemplified by MBT 11, which encompasses regions affected early in AD disease and exhibits high metabolic stress markers. Conversely, primary sensory territories achieve efficiency through molecular streamlining, potentially explaining their relative resilience to age-related decline. Collectively, these molecular profiles provide a context for selective network vulnerability, enabling the prediction of disease risk and suggesting territory-specific therapeutic strategies.

Our bilateral sampling strategy revealed comprehensive proteomic insights into hemispheric specialization. The enrichment of voltage-gated potassium channels (Kv1 family) and bioenergetic components in the left M1 suggests a molecular optimization for high temporal fidelity and fine motor control. Conversely, the right M1’s focus on homeostatic and transcriptional regulators indicates a different computational strategy, perhaps prioritizing circuit stability and inhibitory tone. Additionally, the sex-specific metabolic profiles discovered in the cerebellum, characterized by female prioritization of mitochondrial efficiency via fatty acid metabolism and male preference for glycolytic biogenesis, provide a potential molecular explanation for sex-based differences in susceptibility to metabolic and neuroinflammatory stressors.

The clinical utility of this atlas is demonstrated through analysis of epilepsy-associated proteomic disruptions. By using our healthy proteome as a reference, we identified a thousand consistently dysregulated proteins across epilepsy patients, many of which involve rho family GTPases and axonal guidance. Notably, our network-based prioritization successfully captured SV2A and SCN2B. These established FDA-approved targets validate the clinical precision of our pipeline.

Beyond these markers, the identification of P2RX7 serves as a robust proof of principle for our analytical approach. The upregulation of P2RX7 links seizure activity to neuroinflammation through microglia activation and blood-brain barrier disruption. These findings suggest that targeting the inflammatory-excitatory cycle with P2RX7 inhibitors could provide a superior strategy for refractory epilepsy. This paradigm may set a new standard for mechanism-based drug discovery in neurology.

Despite the unprecedented spatial resolution of this atlas, certain limitations remain. The current study relies on a relatively small cohort of four healthy individuals; while the sampling density is high, future studies incorporating larger cohorts will be necessary to capture the full spectrum of human genetic and environmental diversity. Furthermore, as our analysis utilized bulk tissue punches, the cell-type-specific contributions to these proteomic signatures remain inferred. Integrating this resource with emerging spatial single-cell proteomics will be a vital next step.

In summary, our atlas offers a key map of the brain’s protein landscape. It reveals that brain function relies on specialized protein patterns regulated beyond the genetic level. By mapping the molecules behind brain networks and uncovering new targets for epilepsy, this study helps bridge the gap between brain structure and clinical treatment. The resource will be made freely available through our web portal, enabling researchers worldwide to explore protein expression patterns, compare disease-affected regions, and identify therapeutic opportunities.

## Methods

### Formalin-fixed brain sample collection and MRI imaging

Human brain samples were collected from Dalian Medical University Body and Organ Donation Center. The human brains were formalin-fixed within 10 hours of death. The samples originated from two male and two female donors. Brain punches were taken from the cerebral cortex, subcortical nuclei, and cerebellum. Furthermore, the cerebral cortex samples were segmented into white and grey matter. After sampling, the MRI scanning of the brain was performed to obtain the spatial coordinates of all the sampling locations. The donations are handled in accordance with ethical guidelines and regulations. These samples were then transported to the laboratory for proteomic analysis.

### Proteomic sample processing and data profiling

For proteome data profiling, we employed PCT-assisted sample preparation^19,22,45^ and PulseDIA mass spectrometry data acquisition^23^. PCT utilizes alternating cycles of high pressure and ambient pressure to enhance protein extraction efficiency from small tissue volumes, particularly important for extracting membrane proteins and proteins from compact tissue structures. This method enabled complete protein solubilization from biopsy-level samples (1-2 mg) while maintaining protein integrity. The procedures involved deparaffinization and rehydration of the formalin-fixed brain tissue samples, followed by PCT treatment to enhance protein extraction and digestion efficiency. Then, the samples were desalted. The resulting peptide mixture was analyzed using a hybrid quadrupole-Orbitrap mass spectrometer coupled with a nanoflow-performance liquid chromatography system.

To investigate cross-species differences, we employed FAXP, which combines tissue expansion with protein staining to visualize cytoarchitectural boundaries, allowing accurate punch needle sampling of specific anatomical regions that would be challenging to isolate in unexpanded tissue.

### Hybrid spectral library generation

For spectral library construction, we combined 354 data-dependent acquisition (DDA) runs from fractionated pooled brain tissue with 5362 DIA analyses from individual samples. The DDA runs utilized high-pH reverse-phase fractionation to increase proteome coverage (**Figure S1a**). To generate the hybrid spectral library, we employed MSFragger^46^ and utilized both PulseDIA and DDA files. First, DIA-Umpire generated pseudo-MS/MS spectra from the DIA files. Subsequently, we used MSFragger in DDA mode for the library generation. The default settings were applied, including the generation of decoys and contaminants. Additionally, the ciRT peptide sequences contained in FragPipe were used for retention time normalization. For retention time correction during library generation, the option “ciRT” was utilized. The library was filtered to achieve a 1% false discovery rate at both the protein and peptide levels. The FASTA file, consisting of a reviewed human Swiss-Prot sequence database containing canonical and isoform sequences (51,548 entries), was downloaded from UniProt (https://www.uniprot.org/).

### Proteome and transcriptome integration

We downloaded and processed Allen Brain Atlas microarray data from six neurotypical donors, comprising expression profiles for approximately 20,000 genes across multiple brain regions, to enable the comparison between protein and mRNA expression patterns across matched brain locations^47^. Allen Brain samples were matched to MNI coordinates using their documented stereotactic positions, enabling spatial alignment with our proteomic data. Brainnetome Atlas connectivity matrices were integrated to assign functional network membership to each sample. Data harmonization included gene identifier mapping, batch correction using ComBat^48^.

### Quality Control

Checkpoints Q1–Q3 were established to verify sample accuracy, while Q5–Q11 monitored sample preparation and data acquisition steps to maintain data quality (**Figure S1c**). Batch effects were minimized through stratified sampling process and inclusion of pooled quality control samples every 15 injections (**Figure S1d**).

### Statistical Analysis and Single-Cell Integration

To ensure analytical consistency, both bulk proteomics and single-cell transcriptomic data were standardized into the AnnData format. For scRNA-seq data, only protein-coding genes were retained to harmonize the feature space with proteomic measurements. Differential expression analysis of bulk proteomic data was conducted in Scanpy using the Wilcoxon rank-sum test to compare left and right hemispheres, with differentially expressed proteins (DEPs) defined at p < 0.05. Cell-type-specific molecular signatures were identified using a one-versus-rest Wilcoxon framework, incorporating a downsampling strategy to balance group sizes between target cell types and controls. Owing to the high statistical power of large-scale single-cell datasets, marker genes were defined as the top 300 features per cell type ranked by test statistics.

More details are provided in Supplementary Methods.

## Data Availability

The mass spectrometry proteomics data generated in this study have been deposited in the iProX repository under the accession number PSV023 and are available at https://www.iprox.cn/page/PSV023.html;?url=1774082320375mAUj

password: wNmV

In addition, all processed data and associated resources are freely accessible through the ProtTalks platform at https://db.prottalks.com

## Acknowledgments

We would like to express our gratitude to the National Key R&D Program of China (Grant No. 92259201, 2023C03056 and 2021ZD0200200). We are also grateful to the patients participated in this project for their donation of body or body tissue for scientific research. We also appreciate the advanced computing services provided by Westlake University High-Performance Computing Center.

## Author contributions

Conceptualization: T. G., Y. L., T. J., and Y. M. Methodology: T. G., T. J. and Y. L. Investigation: Q. X., Y. X., M. L., Z. Y., J. Gao., J. Y., W. J., and W. G. Resources: Y. L., Y. M., L. C., H. Y., M. L., X. Z., N. F., and F. X. Data analysis: Q. X., Y. X., Z. Y., J. Y., W. J., W. G., J. L., W. L., and J. G. Visualization: Q. X., Y. X., Z. N., Y. C., and Z. F. Writing-origin draft: Q. X., Y. X., and J. Gao. Writing-review & editing: T. G., T. J., E. S. L., Z. Y., and Y. S.

## Competing interests

T. G. is a shareholder of Westlake Omics Inc. The remaining authors declare no competing interests.

## Additional information

Supplementary Information is available for this paper. Correspondence and requests for materials should be addressed to Tiannan Guo.

## Supplementary figure legend

**Figure S1.**
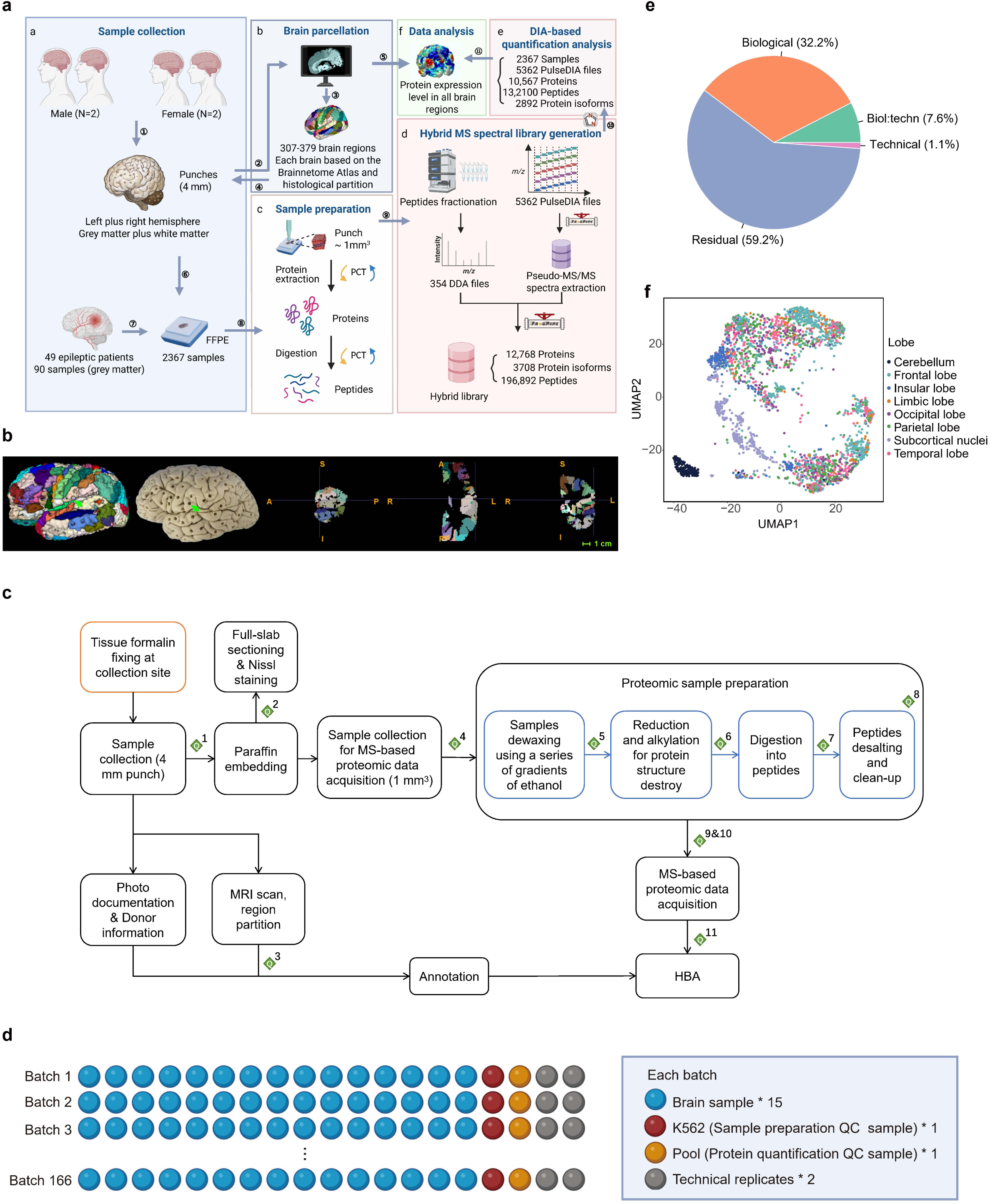
Study design and quality control workflow, related to Figure 1. **a,** Comprehensive data generation and analysis pipeline overview. Brain tissue collection encompassed four donors (two male, two female), with 307-379 tissue punches collected per brain (total n = 2367 samples). Cerebral cortex samples were anatomically separated into white and grey matter prior to paraffin embedding. Additionally, 90 epileptic brain grey matter samples were collected for comparative proteomic analysis. MRI scanning was performed to obtain precise spatial coordinates for all sampling locations of four donor brains, which were subsequently mapped to the Brainnetome Atlas reference framework. Paraffin-embedded specimens were processed to obtain 0.6-1.0 mg samples for PCT-assisted protein extraction and enzymatic digestion. The mass spectrometry workflow employed a DDA-DIA hybrid approach: DIA-Umpire generated pseudo-MS/MS spectra from DIA files, and DIA-NN executed library-based peptide identification and quantification. **b,** The 3D brain reconstruction shows sampling locations (indicated by green arrows) that correspond exactly to anatomical positions in the accompanying photograph. The three orthogonal MRI views (sagittal, horizontal, and coronal planes) demonstrate the spatial precision achieved for each sampling point, enabling accurate anatomical assignment and coordinate mapping. **c,** Quality control checkpoint workflow integrated throughout the experimental pipeline. Green diamond symbols (Q) indicate mandatory quality control steps implemented between major processing stages. Comprehensive quality control procedures are detailed in Supplementary Methods 1, Section VI (Quality Control), covering sample collection, processing, mass spectrometry acquisition, and data analysis validation steps. **d,** Batch design strategy for systematic brain sample analysis to minimize technical variation. Each analytical batch contained: one pooled quality control sample, 15 brain tissue samples, one human cell line sample for process monitoring, and two technical replicates, ensuring balanced representation and robust quality assessment. **e,** Dataset variance decomposition analysis using principal component analysis. The pie chart illustrates the relative contributions of different variance sources: biological variables (brain region, sex, individual differences). **f,** Uniform Manifold Approximation and Projection dimensionality reduction visualization of the complete proteomic dataset. Each point represents a single brain sample.

**Figure S2.**
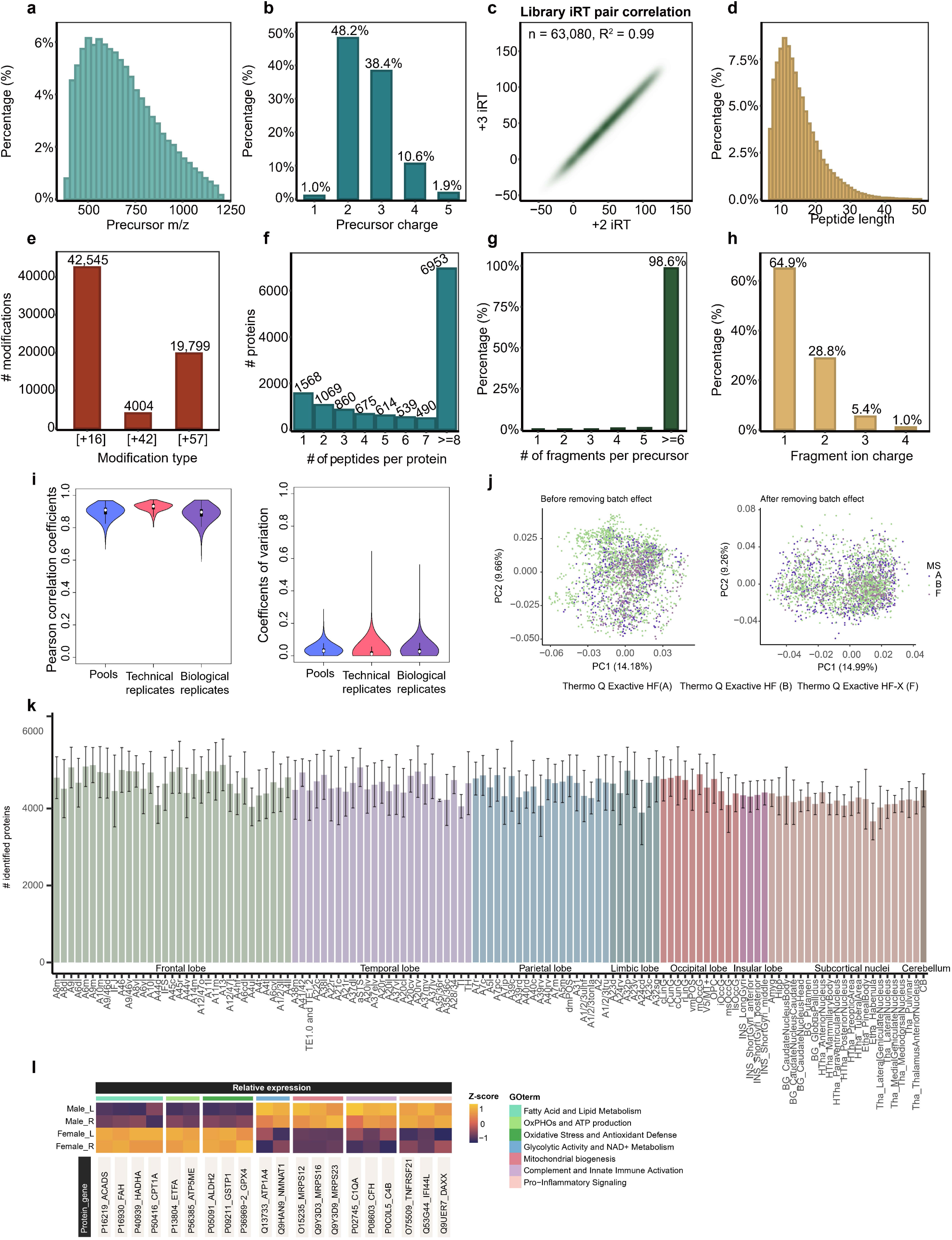
Characterization of the spectral library of human brain and quatification of proteome data, related to Figure 1. **a,** Mass-to-charge ratio (m/z) distribution of peptide precursors in spectral library. **b,** Distribution of precursor ion charge states detected in the spectral library. **c,** Quality assessment of retention time alignment using indexed retention time (iRT) correction. **d,** Distribution of identified peptide lengths in the spectral library. **e,** Frequency analysis of post-translational modifications detected in the dataset. Bar chart displays counts for three major modifications: methionine oxidation (Δmass +16 Da), N-terminal acetylation (Δmass +42 Da), and cysteine carbamidomethylation (Δmass +57 Da. **f,** Distribution of proteotypic peptides per protein in the spectral library. **g,** Fragment ion coverage analysis showing the number of fragment ions per precursor. **h,** Charge state distribution of fragment ions observed in MS/MS spectra. **i,** Reproducibility assessment across different sample types. Left panel shows Pearson correlation coefficients; right panel displays coefficients of variation (CV) for quality control pooled samples, technical replicates, and biological replicates. **j,** Principal component analysis of dataset variance across three mass spectrometry instruments. Scatter plots show PCA clustering before (left) and after (right) batch effect correction, with colors representing different instruments. **k,** Protein quantification coverage across all brain subregions. Bar chart displays the number of quantified proteins in each anatomical region. Colors represent different brain divisions: FL, TL, PL, LL, OL, IL, subcortical nuclei, and cerebellum. **l,** Heatmap displaying protein expression patterns for significantly dysregulated proteins in the cerebellum. Color scale represents z-score normalized protein expression levels, with orange indicating higher expression and dark purple indicating lower.

**Figure S3.**
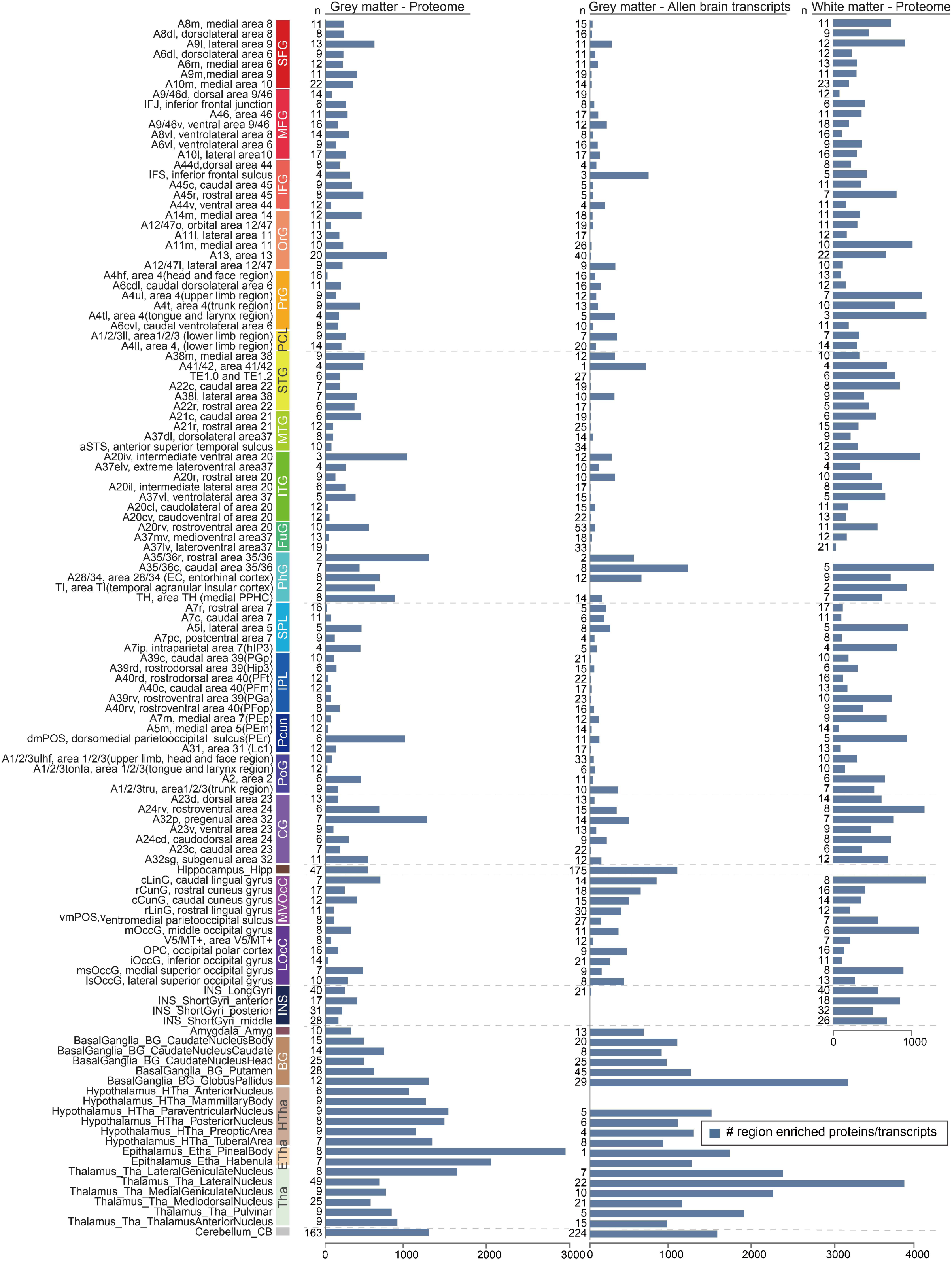
RE proteins and transcripts distribution across brain subregions, related to Figure 2. Regional distribution of RE features across anatomically defined brain subregions. Left panel: number of RE proteins enriched in grey matter across 121 subregions, using specificity score > 0.93 (95th percentile threshold of the protein expression matrix) and ANOVA with Benjamini-Hochberg adjusted p < 0.05. Middle panel: number of RE transcripts enriched in grey matter across the same 121 subregions, based on microarray expression data from the Allen Human Brain Atlas, using specificity score > 0.94 (95th percentile threshold of the transcript expression matrix) and ANOVA with Benjamini-Hochberg adjusted p < 0.05. Right panel: number of RE proteins enriched in white matter across 94 subregions, applying identical protein enrichment criteria.

**Figure S4.**
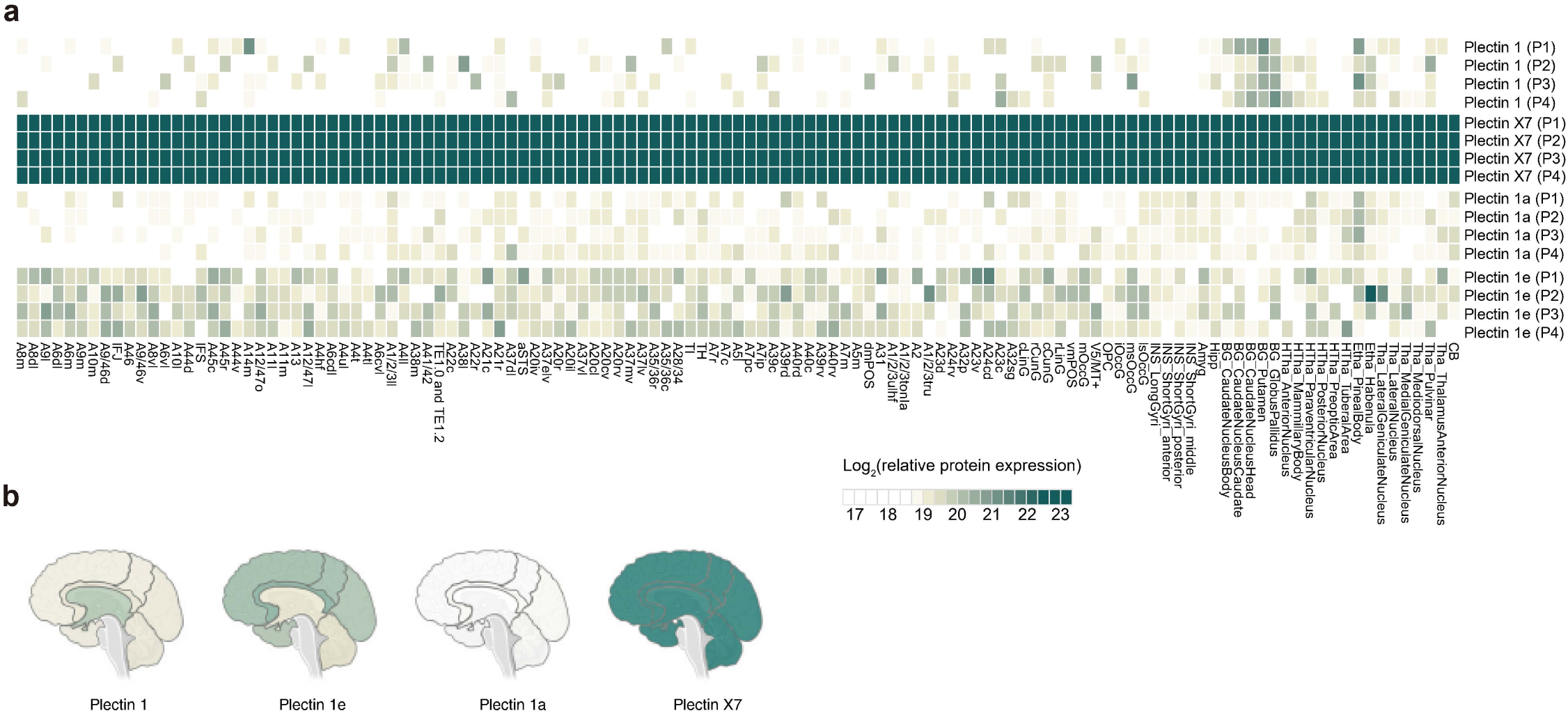
Plectin protein isoforms, related to Figure 1. **a,** Regional expression analysis of Plectin protein isoforms across brain subregions. Four Plectin isoforms (Plectin 1, Plectin 1e, Plectin 1a, and Plectin X7) were quantified across 121 anatomically defined subregions. **b,** Brain region heatmap displaying mean expression levels of the four Plectin isoforms across major anatomical divisions: FL, TL, PL, LL, OL, IL, subcortical nuclei, and cerebellum. Color intensity represents relative expression levels.

**Figure S5.**
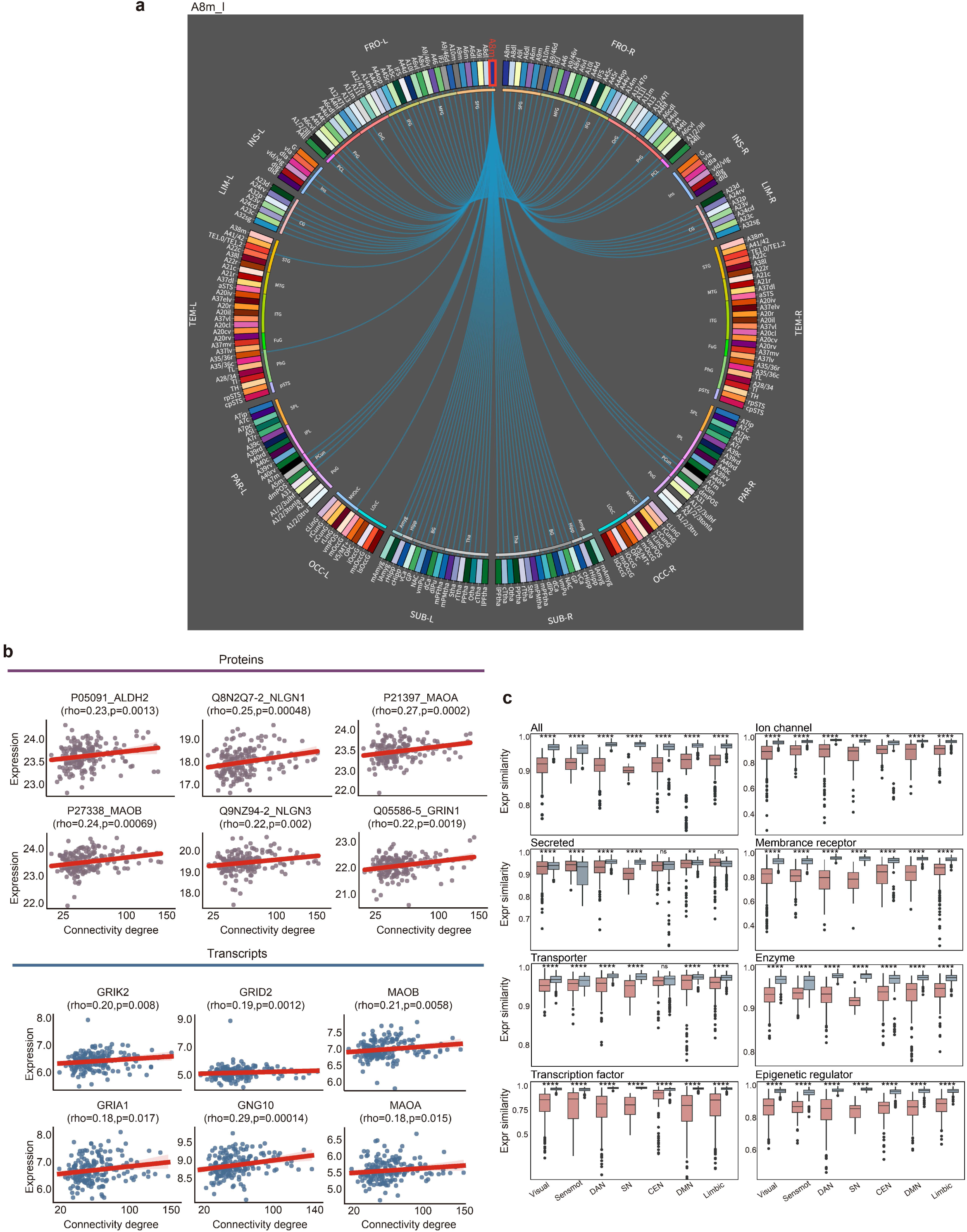
Structural connectivity patterns visualized through connectograms, related to Figure 3. **a,** Connectogram displaying structural connectivity patterns for the left hemisphere area A8m (medial frontal area 8). Connecting lines represent strong structural connections between left A8m and other brain subregions, based on diffusion tensor imaging tractography data from the Brainnetome connectome atlas. Complete connectograms for all brain regions analyzed in this study are available through the Brainnetome Atlas website (https://atlas.brainnetome.org/). **b,** Scatter plots demonstrating the correlation between structural connectivity degree and protein expression levels across brain subregions. Connectivity degree calculated as the sum of strongly connected subregions for each given subregion. Each point represents a single subregion. Spearman correlation coefficients (rho) and Benjamini-Hochberg adjusted p-values are indicated at the top of each panel. **c,** Scatter plots showing the correlation between structural connectivity degree and transcript expression levels across brain subregions. **c,** Box plots comparing correlation strengths between highly connected brain subregion pairs across different gene functional categories. Proteins and transcripts were classified into eight functional groups: All, Ion channel, Secreted, Membrane receptor, Transporter, Enzyme, Transcription factor, Epigenetic regulator.

**Figure S6.**
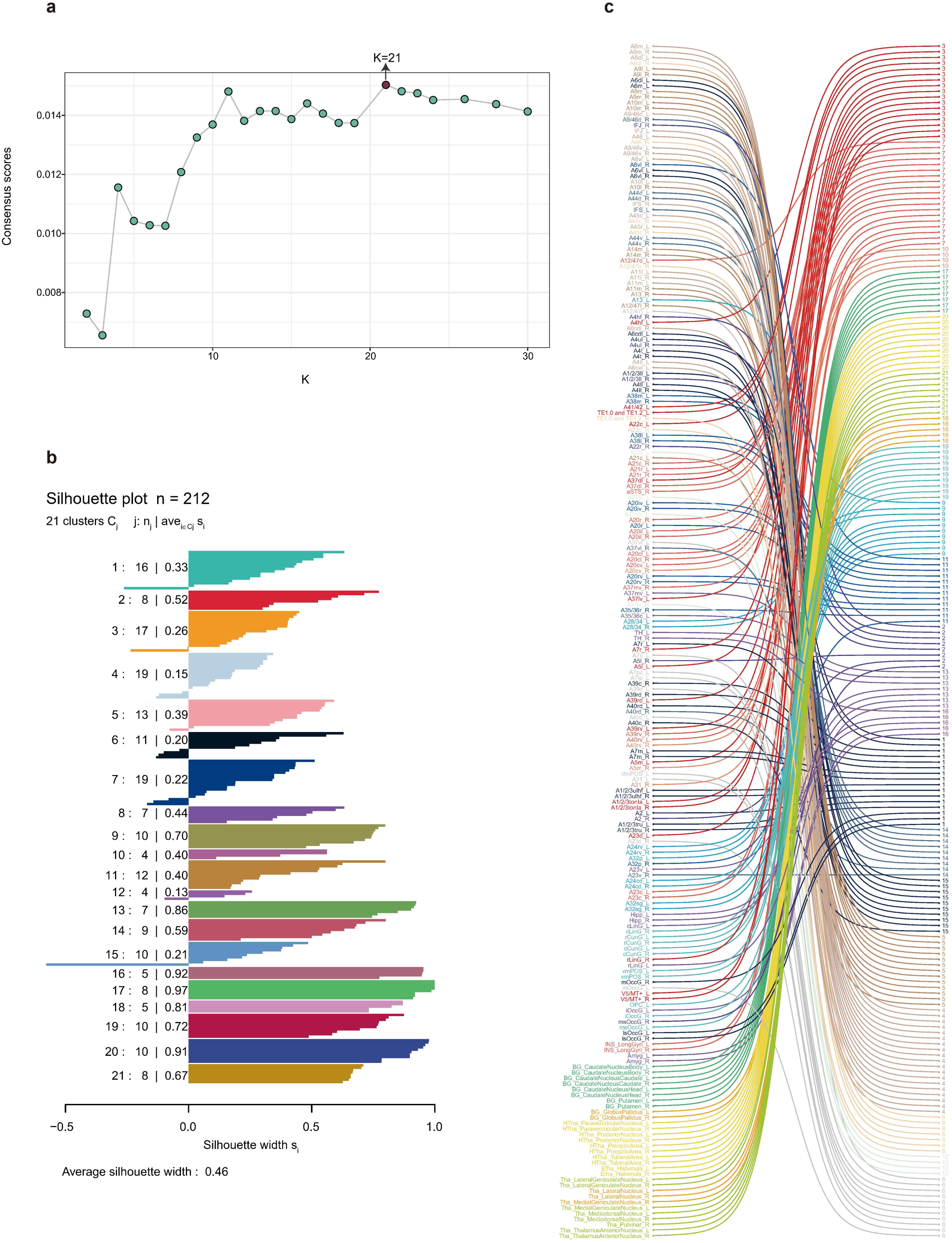
Molecular-based brain parcellation, related to Figure 4. **a,** Consensus clustering optimization analysis showing the relationship between consensus scores and cluster number (K) across a range of K=1 to 45. **b,** Silhouette analysis validation of the 21-cluster solution. The silhouette plot demonstrates cluster quality with an average silhouette width of 0.46. Each bar represents an individual brain subregion, with positive values indicating good cluster assignment and negative values suggesting potential misclassification. **c,** Cluster composition and subregion assignment overview. Line plot connecting subregion names (left) to their assigned molecular clusters (right, numbered 1-21).

**Figure S7.**
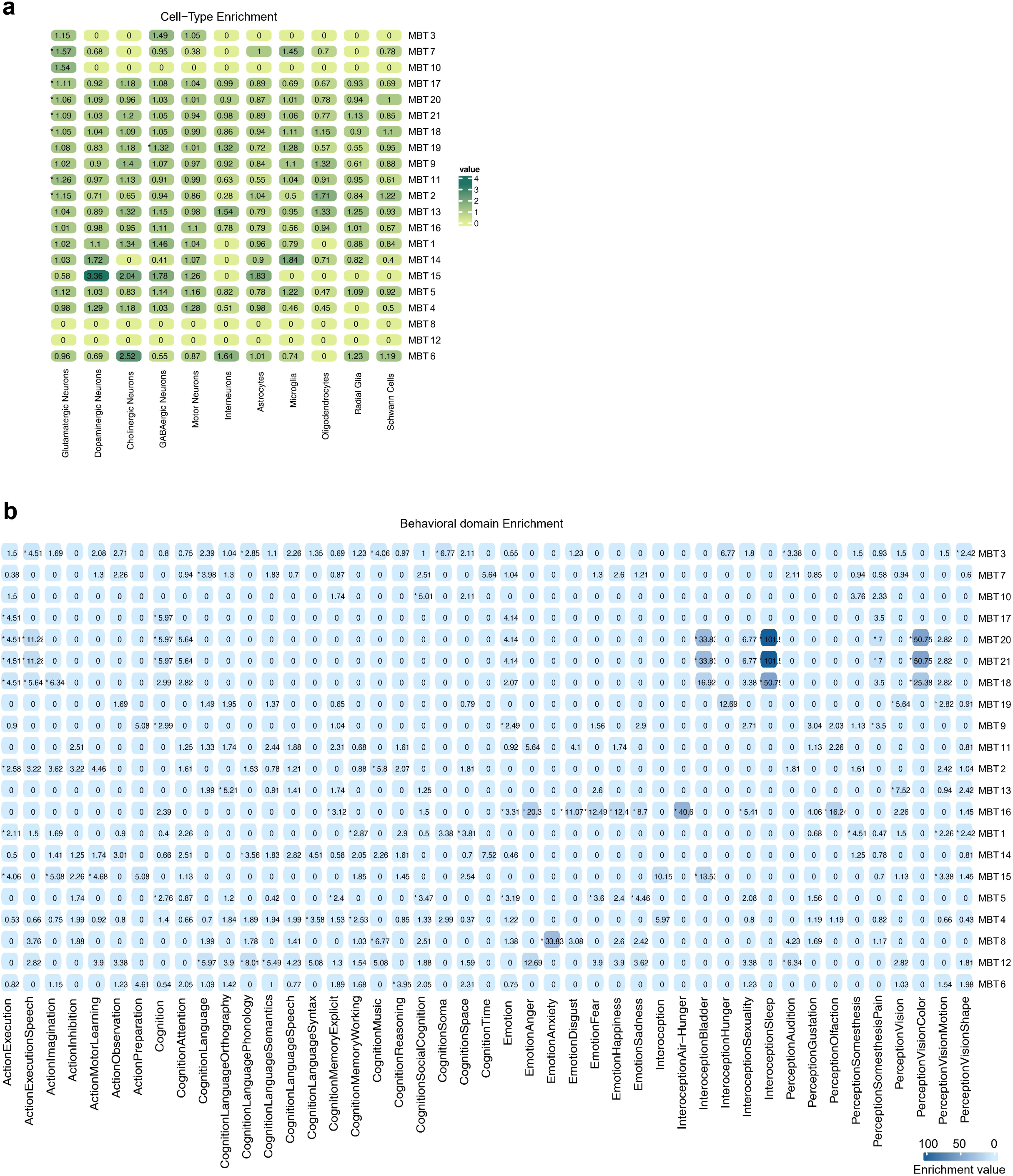
Neuronal signalling, development, and support pathway analysis, related to Figure 4. **a,** Cell-type enrichment analysis for each of the 21 molecular clusters using established cell-type markers from the Human Protein Atlas (HPA). Enrichment scores were calculated using clusterProfiler. The heatmap shows enrichment scores for different cell types across clusters, with asterisks (*) marking statistically significant enrichments (p < 0.05, Fisher’s exact test). **b,** Functional behavioral domain enrichment analysis across the 21 molecular clusters. Ridge plot shows the distribution of enrichment scores (calculated using clusterProfiler) for each cluster across 42 behavioral domain profiles from the BrainMap database (brainmap.org)^37^. Enrichment significance was determined using Fisher’s exact test with p < 0.05.

**Figure S8.**
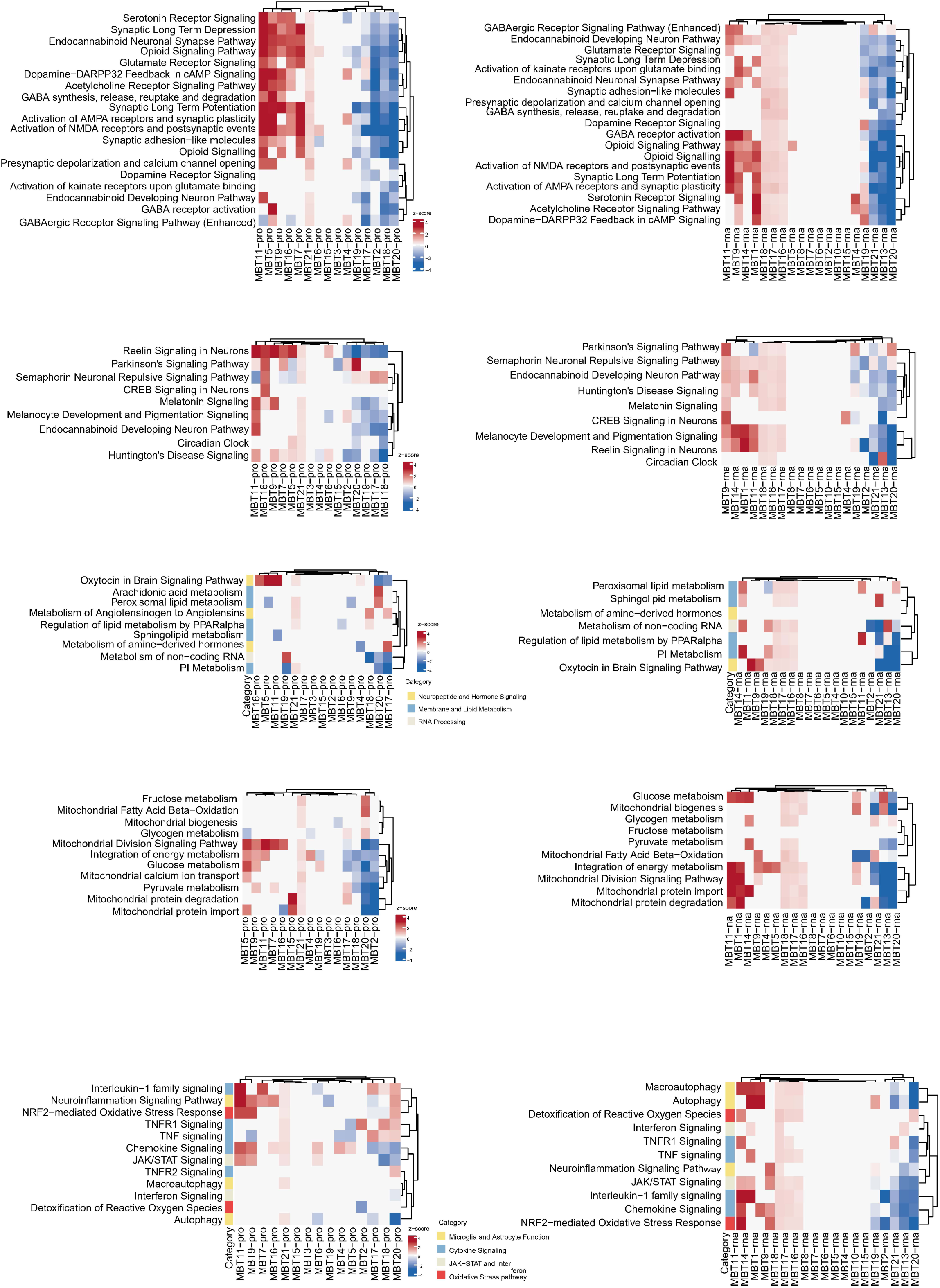
Functional pathway analysis reveals territory-specific biological programs, related to Figure 4. Ingenuity Pathway Analysis (IPA) activation z-scores for major biological themes across MBTs. Pathways are grouped into five functional categories: neurotransmitter signaling, brain development, energy metabolism, lipid metabolism, and immune response. Red indicates pathway activation (positive z-score) and blue indicates suppression (negative z-score).

**Figure S9.**
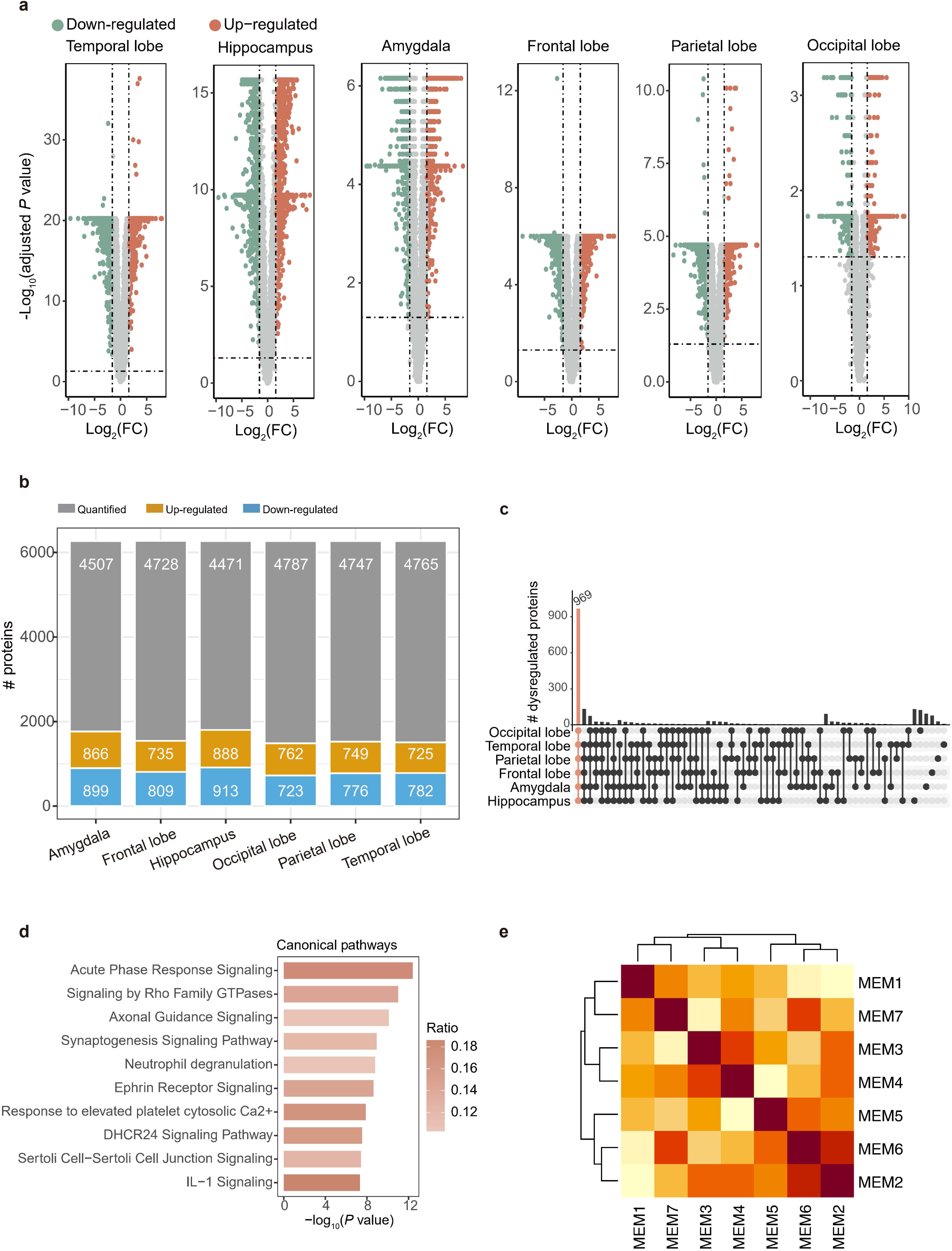
Differential protein analysis between epileptic and healthy brains across six brain regions, related to Figure 5. **a,** Volcano plots displaying dysregulated proteins in six brain regions (Temporal lobe, Hippocampus, Amygdala, Frontal lobe, Parietal lobe, Occipital lobe). Points outside the significance thresholds (|Fold change| > 3, Benjamini-Hochberg adjusted p < 0.05, Wilcoxon test) represent dysregulated proteins in epileptic versus healthy brain tissue. **b,** Bar plot quantifying proteomic coverage and dysregulation across the six brain regions. For each region, bars show the total number of proteins quantified (grey bars) and the subset of significantly dysregulated proteins in epileptic samples. **c,** UpSet diagram illustrating the overlap of dysregulated proteins across the six brain regions. **d,** Canonical pathway enrichment analysis of the 969 dysregulated proteins shared across all six brain regions using IPA.

**Supplementary Figure S10.**
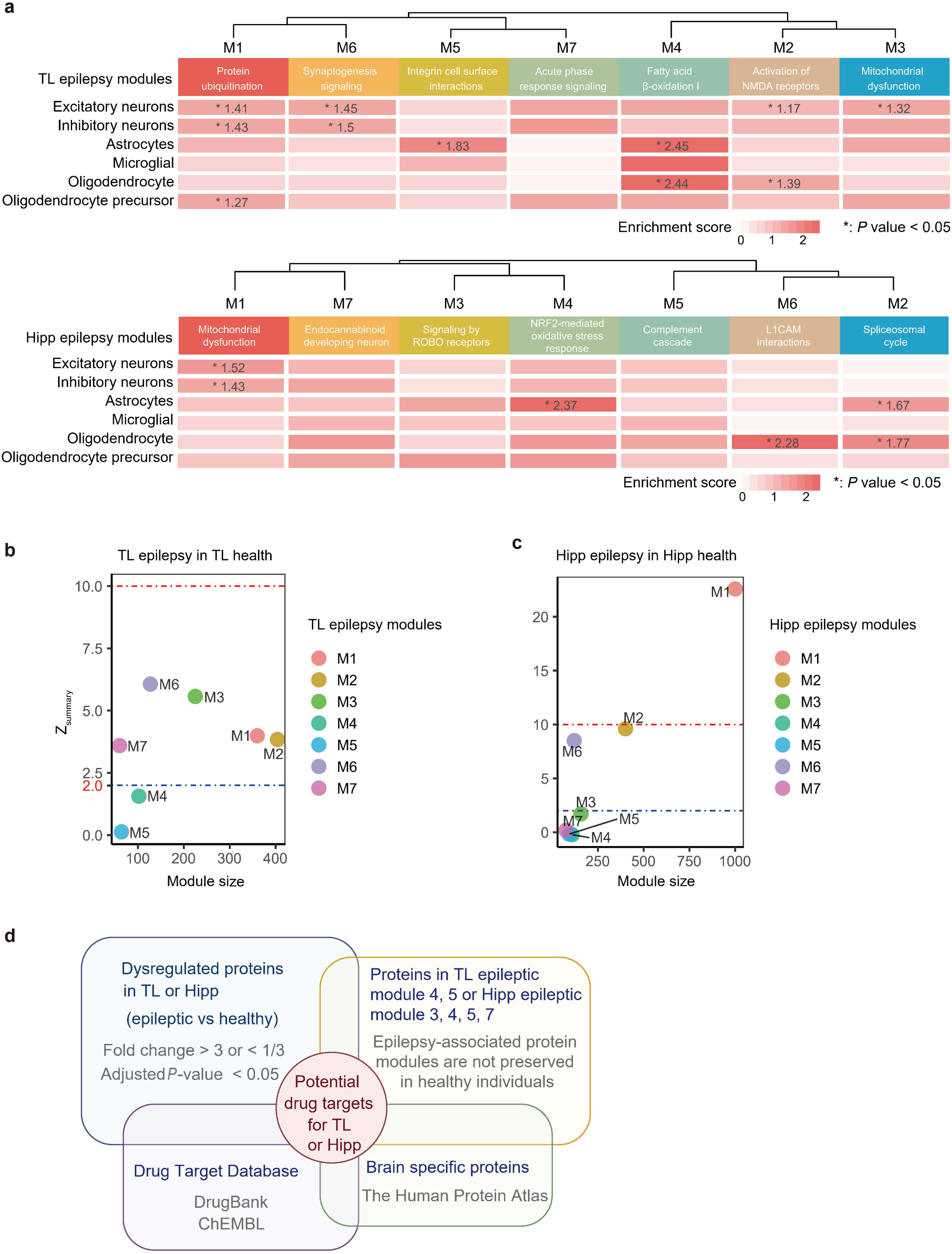
Anti-epilepsy drug target identification through co-expression network analysis, related to Figure 5. **a,** WGCNA of epileptic temporal lobe samples revealing seven distinct protein co-expression modules. Each module represents a group of proteins with highly correlated expression patterns. Canonical pathway analysis using IPA identified the biological processes associated with each module. Cell-type specificity was assessed through enrichment analysis using neuron- and glia-specific protein markers from the HPA. **b,** Module preservation analysis comparing epileptic TL protein networks with healthy TL networks. Modules with preservation Z-summary scores < 2 were considered non-preserved (indicating epilepsy-specific disruption). Modules with Z-summary scores ≥ 2 (blue dotted line) were considered preserved, while those with scores ≥ 10 (red dotted line) were highly preserved. X-axis represents the number of proteins in each module. **c,** Module preservation analysis comparing epileptic temporal lobe protein networks with epileptic hippocampus networks. This cross-regional comparison identifies modules that are specifically disrupted in temporal lobe epilepsy versus those representing general epilepsy-associated changes. Preservation thresholds are identical to panel b: Z-summary < 2 (non-preserved), ≥ 2 (preserved, blue line), and ≥ 10 (highly preserved, red line). **d,** Systematic drug target selection criteria for epileptic temporal lobe therapy.

**Supplementary Figure S11.**
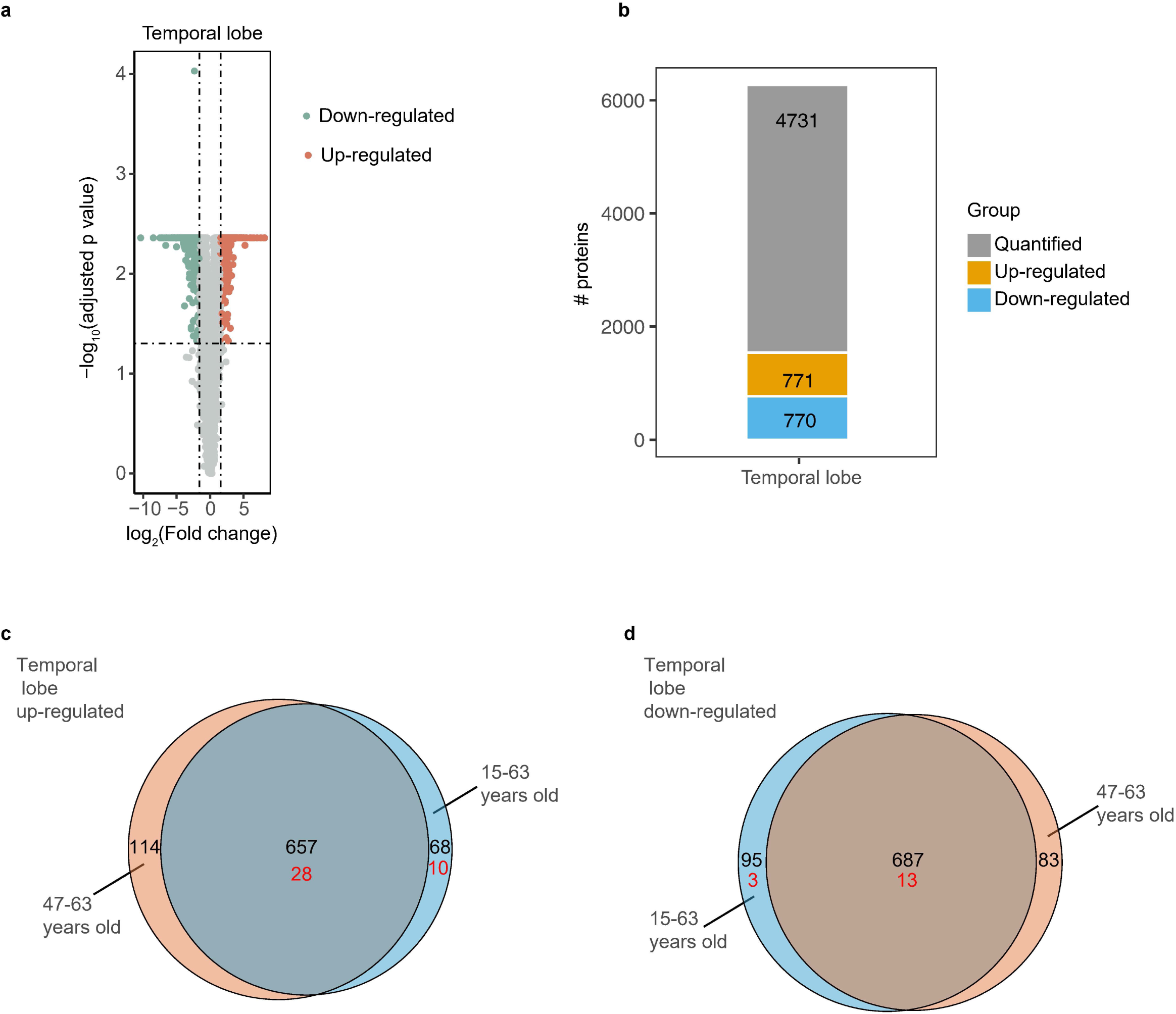
Evaluation of age-related confounding effects on epilepsy-associated proteomic alterations in the temporal lobe, related to Figure 5. **a,** Differential expression analysis of the age-matched temporal lobe samples (n=4 pairs, aged 47–63 years). Differentially expressed proteins (DEPs) (Fold change > 3, adjusted p value< 0.05, Wilcoxon signed-rank test) are highlighted in orange (up-regulated) and green (down-regulated). **b,** Quantification of total and DEPs in the age-matched analysis. **c and d,** Venn diagrams showing the conservation of up-regulated (**c**) and down-regulated (**d**) proteins between the age-matched group and the full cohort (15–63 years). Red numbers represent identified drug targets.

## Notes

### Summary of Updates

Author affiliations updated, Figure 1 and Figure 2 revised, supplemental files uploaded.

