## Supplementary information for "A spatially resolved human brain proteome atlas for decoding brain function and disease"

#### Supplementary Methods

##### 1. Human brain sample collection and processing

##### 1.1 Donor selection criteria and demographics

Human postmortem brain samples (n = 4) were obtained from the Body and Organ Donation Center of Dalian Medical University (see Table 1 for donor information). All donors had voluntarily consented to body donation for scientific research prior to death, or consent was provided by their legal representatives. None of the donors had a history of neurological or neurodegenerative disorders. The human bodies were formalin-fixed within 10 hours of death to ensure proper preservation of the brain tissue. The brains were extracted from the skull and then immersed in formalin for at least four weeks for adequate fixation. All procedures were conducted in accordance with institutional protocols and ethical regulations governing the use of human tissues for research.

Table 1 Donator information

| **Individual** | **Sex** | **Age** | **Postmortem interval** |
|---|---|---|---|
| Healthy P1 | Male | 62 | 8 hours |
| Healthy P2 | Female | 84 | 4 hours |
| Healthy P3 | Male | 67 | 7 hours |
| Healthy P4 | Female | 51 | 8 hours |

Collecting the postmortem brain samples was conducted in compliance with all relevant regulations and ethical guidelines, and informed consent was obtained from the donors or their legal representatives before donation. The samples were obtained to advance scientific research and our knowledge of neuroscience.

##### 1.2 Brain dissection protocols

Postmortem brains were dissected using standardized neuroanatomical procedures to ensure spatial consistency across donors. The skull was opened circumferentially with an autopsy saw, and the calvarium was removed together with the attached dura mater. The brain was carefully extracted by releasing the posterior fossa contents through dissection of the tentorium margins. Following removal, the intact brain was weighed, visually inspected, and photographed from multiple angles to document gross anatomy. The brain was bisected along the mid-sagittal plane using a long knife, beginning at the brainstem and proceeding through the cerebellar vermis, diencephalon, and corpus callosum. Throughout dissection, spatial orientation was preserved, and photographic documentation was carried out at each stage to facilitate subsequent MRI-based spatial registration. All dissections were conducted by trained personnel using consistent anatomical landmarks and tools to minimize inter-sample variability.

#### 1.3 Formalin fixation procedures

The four postmortem brain samples were immersed in 10% formalin within 10 hours postmortem. Brains remained immersed for a minimum of 14 days without agitation, with containers gently inverted daily to promote uniform fixation. After fixation, specimens were transferred into a 0.9% saline solution containing 0.05% sodium azide to inhibit microbial growth and stored at 4 °C until sample collection and imaging. This protocol was optimized to preserve both tissue morphology and protein integrity, enabling high-quality ex vivo MRI and downstream proteomic analysis. Prior to scanning, brains were placed in a custom-designed, MRI-compatible holder that stabilized the specimen and preserved anatomical orientation throughout the imaging process.

#### 1.4 Tissue punching methodology and spatial registration

Sampling targeted the cerebral cortex, subcortical nuclei, and cerebellum, using a sterile stainless-steel punch needle with a 4 mm diameter. Each brain yielded 307-379 spatially distributed locations, systematically sampled to ensure broad anatomical coverage. Cortical punches were further classified into gray and white matter based on gross anatomical features and myelination contrast. A total of 541-608 tissue samples were obtained for each brain. Throughout the process, anatomical orientation was maintained using surface photographs and dissection maps, enabling accurate spatial registration. Sampling coordinates were recorded and cross-referenced with ex vivo MRI images to facilitate downstream spatial mapping and integrative analysis.

#### 1.5 Sample storage and quality control

Following dissection, all tissue samples were stored at –80 °C for long-term preservation. Freezer temperatures were continuously monitored using automated logging systems with alarm-triggered notifications for deviations. Each sample was barcoded and tracked using a laboratory information management system linking sample ID, spatial coordinates, brain region, and donor metadata. Quality control (QC) was performed at multiple stages, including visual inspection for tissue integrity and protein concentration quantification using BCA assay. Periodic reassessment of archived samples was conducted to confirm protein stability over time. Spatial registration data and QC records were integrated into the database to ensure traceability and support reproducible downstream analysis.

#### 1.6 Ethical approvals

All procedures involving human and animal tissues were approved by the relevant institutional ethics committees and conducted in accordance with the Declaration of Helsinki. Human brain tissues from neurologically healthy donors were obtained through the Brain Donation Centers at Dalian Medical University (Ethics Approval ID: 201905) and Shanghai Jiao Tong University (Ethics Approval ID: 2022-A-01). Informed consent for brain donation and research use was obtained from all donors or their legal representatives prior to death. Epilepsy patient brain tissues were collected during clinically indicated surgical procedures at Huashan Hospital with approval from the Huashan Hospital Ethics Committee (Ethics Approval ID: KY2022-590. Mouse brain collection procedures were approved by the Institutional Animal Care and Use Committee (IACUC Approval ID: AP#25-027-GTN). Sample storage, processing, and proteomic analysis were conducted at Westlake University under Ethics Approval ID: 20240527GTN002. All tissue handling, storage, and distribution followed established protocols for maintaining donor anonymity while enabling scientific reproducibility.

#### 2. MRI imaging and spatial mapping

##### 2.1 MRI data acquisition

All brain samples were scanned using a 1.5 Tesla MRI system equipped with a 32-channel head coil. All four brain parts were positioned within a single scan to preserve their relative spatial orientation.

##### 2.2 Image preprocessing and quality control

The acquired MRI data were processed using a combination of ANTs (v2.4.3) and SPM12 software packages. Preprocessing included intensity inhomogeneity correction using the N4 algorithm, followed by brain tissue segmentation and subcortical structure segmentation.

Quality control was performed throughout the pipeline to ensure data accuracy and reproducibility. This included routine scanner calibration, evaluation of signal-to-noise ratio, and visual inspection of segmentation results and anatomical structures.

##### 2.3 Brainnetome Atlas registration

Following preprocessing, the reconstructed brain image was divided into left and right hemispheres to account for potential misalignment introduced during sample preparation. Each hemisphere was independently registered to the corresponding hemisphere of the Brainnetome Atlas.

Initial alignment was performed using a 6-degree-of-freedom (6 DOF) rigid transformation, followed by nonlinear registration to refine anatomical correspondence. The resulting deformation field was applied to the Brainnetome parcellation to generate atlas labels in individual brain space.

##### 2.4 Sample localization and annotation

Sample localization and anatomical annotation were performed using a multi-step procedure:

(1) Surface reconstruction. A surface mesh was generated from the segmented brain volume using the Marching Cubes algorithm, enabling visualization of tissue morphology and sampling sites.

(2) Punch hole identification. Sampling sites (punch holes) were identified through visual inspection of the reconstructed surface. The voxel-space coordinates of each hole center were recorded.

(3) Sample labeling. Each punch hole was assigned a unique sample ID by integrating voxel coordinate information with photographic records obtained during sample collection.

(4) Atlas-based annotation. After mapping the Brainnetome Atlas into individual brain space, the recorded coordinates of each sample were used to determine its corresponding anatomical region based on atlas labels.

(5) Quality validation. To ensure annotation accuracy, all assignments were manually verified by overlaying punch hole locations with atlas regions on the reconstructed brain surface.

This pipeline enables precise spatial correspondence between proteomic samples and anatomically defined brain regions.

#### 3. Proteomic analysis

##### 3.1 Batch design for brain sample analysis

We used a stratified sampling method to minimize batch effects in our brain sample analysis to distribute 2291 brain samples into 166 batches. The factors considered for stratification included hemisphere, white or gray matter, individual, and brain region. Each batch contained one quality control pool sample, 15 brain samples, one human cell line sample for monitoring the sample preparation process, and two technical replicates (**Supplementary Fig. 1d**).

The 15 brain samples in each batch were chosen based on the stratified factors. The quality control pool sample was created by pooling equal amounts of proteins from each brain sample in the batch and served as a reference for assessing batch effects and technical variability. The two technical replicates were included to determine the reproducibility of the analysis.

The batch design was carried out using ProteomeExpert[1]. The samples in each batch are as similar as possible concerning the stratification factors while minimizing the potential for batch effects. Including reference samples and technical replicates further ensured the accuracy and reproducibility of our results.

##### 3.2 PCT-assisted sample preparation of brain FFPE samples.

The sample preparation process was roughly the same as described previously[2,3]. In brief, each FFPE specimen was punched to obtain 0.6-1.0 mg samples, which were dewaxed using heptane and then hydrated with gradient alcohol. Then, the samples were washed twice in 100% ethanol for 5 minutes and twice in 70% ethanol for 5 minutes. Acid hydrolysis was performed using 0.1% formic acid, and the product was placed in a PCT microtube for base hydrolysis with Tris-HCl (pH=10, freshly prepared) at 95°C for 30 mins to reverse the formaldehyde cross-linking followed by quick cooling. A lysis buffer (6M urea, 2M thiourea) was added along with Tris (2-carboxyethyl) phosphine (TECP) for reduction and iodoacetamide for alkylating the cysteine residues.

PCT-assisted lysis was performed using the Barocycler model NEP2320-Enhanced (Pressure Biosciences Inc., MA) with 90 cycles, 45k psi, 30s HP, 10s AP, and 30°C. We added lysC (enzyme:substrate concentration ratio is 1:80, MS grade, Hualishi Tech. Ltd) and trypsin (enzyme:substrate concentration ratio is 1:20, sequencing grade modified, Hualishi Tech. Ltd) for digestion using the PCT system under the following conditions: 120 cycles, 20k psi, 50s HP, 10s AP, and 30°C. The digestion process was terminated with 15% formic acid. As previously described, the peptides were desalted and cleaned up using a C18 cartridge (Thermo Fisher Scientific™, San Jose, USA). Protein concentration was measured using a NanoDrop™ Nano-500 spectrophotometer at 280 nm using 1-2 µL lysate. Measurements were performed in triplicate with lysis buffer as the blank control.

##### 3.3 Sample preparation for library building.

The peptide sample from each fraction for library generation was resuspended in MS buffer (0.3 µg/µl using 0.1% formic acid and 2% acetonitrile in MS-grade water. Samples from each brain lobe were mixed into a pooled sample according to the different partitions, which included the frontal lobe, temporal lobe, parietal lobe, occipital lobe, limbic lobe, and subcortical nuclei. We use 400 μg peptides for high-pH fractionation, and 200 μg for strong cation exchange Chromatography (SCX) fractionation From each lobe. The mixed samples were dried by vacuum centrifugation (CentriVap, Labconco, Kansas City, USA) at 45°C.

The pooled peptides from each lobe were resuspended in HPLC-grade water at pH=10. Fractions were performed using a Thermo Ultimate 3000 LC system (Thermo Fisher Scientific™, San Jose, USA). The separation was performed using an HPLC C18 column (XBridge BEH, diameter 4.6 mm, length 25 cm, 5 μm particle size). The peptide sample was loaded onto the column and separated using a linear gradient of buffer B (98% acetonitrile, pH = 10) from 5% to 95% over 120 minutes at a 1 mL/min flow rate. We collected 120 fractions at 1-minute intervals using the fraction collector. We set the fraction collector to collect fractions into 96-well plates and programmed the system to collect one fraction per well. We combined the 120 fractions into 30 fractions by pooling every fourth fraction. For example, we combined fractions 1, 31, 61, and 91 into fraction 1; we combined fractions 2, 32, 62, and 92 into fraction 2; and so on until all 120 fractions were combined into 30. We dried the combined fractions using a vacuum concentrator and resuspended the peptides in MS buffer.

The dried peptide pool from each lobe was resuspended in SCX buffer (25% acetonitrile, five mM $KH_2PO_4$, and formic acid to reach pH = 3.0). Next, the sample was loaded into the prepared SCX Cartridge (The Nest Group, BioPureSPN). Then, the peptides were eluted by a series of gradients of buffer B (25% acetonitrile, ten mM $KH_2PO_4$, 500 mM KCl, and formic acid to reach pH=3). The concentrations of KCl in the elution buffers were 50 mM, 100 mM, 150 mM, 250 mM, and 350 mM. Next, peptide desalting and clean-up were performed using a C18 cartridge (Thermo Fisher Scientific™, San Jose, USA). Finally, we dried the combined fractions using a vacuum concentrator and resuspended the peptides in MS buffer.

#### 3.4 Mass spectrometry analysis

##### 3.4.1 DDA-MS

The peptide sample from each fraction, after being combined, were analyzed using a Q Exactive HF-X mass spectrometer (Q Exactive HF-X hybrid quadrupole-Orbitrap mass spectrometer, Thermo Fisher Scientific™, San Jose, USA) and a Q Exactive HF mass spectrometer (Q Exactive HF hybrid quadrupole-Orbitrap mass spectrometer, Thermo Fisher Scientific™, San Jose, USA) coupled to a nanoflow high-performance LC system (DIONEX UltiMate 3000 System, Thermo Fisher Scientific™, San Jose, USA). For the DDA data acquisition, the peptides were separated by a nanoEasy-nLC 1200 System (pre-column: 3 μm, 100 Å, C18 20 mm*75 μm i.d.; analytical column: 1.9 μm, 120 Å, C18 150 mm*75 μm i.d.; Thermo Fisher Scientific) coupled to a Q Exactive HF-X hybrid Quadrupole-Orbitrap mass spectrometer (Thermo Fisher Scientific). For 354 DDA files, the peptides were separated by a 60-min LC gradient (from 3% to 28% buffer B; buffer A: 98% H2O, 2% ACN, 0.1% FA; buffer B: 98% ACN, 2% H2O, 0.1% FA). A full MS scan ranged from 390 m/z to 1,210 m/z with a resolution of 60,000 in the Orbitrap using an Automatic Gain Control (AGC) target of 3e6 charges and a maximum ion injection time of 80 ms. Then, the top 20 precursors from each MS scan were selected to be fragmented and measured at a resolution of 30,000 with an AGC target of 1e5 and a maximum ion injection time of 100 ms.

##### 3.4.2 DIA-MS using multi-injection pulsed gas-phase fractionation

DIA is a proteomic method that quantifies hundreds to thousands of proteins in a single experiment. PulseDIA is a method based on DIA that our lab released, which could improve the coverage of the proteome by equally dividing the conventional DIA analysis covering the entire mass range into multiple injections with complementary windows[4]. We employed two parts: 30 min gradient PulseDIA method combined on Q Exactive HF-X and Q Exactive HF mass spectrometer.

To generate PulseDIA files for 2291 brain samples, 500 ng peptides were injected twice using a 30-min LC gradient (3% to 28% buffer B). The m/z range of MS1 was 390 to 1210 at a resolution of 60,000, with an AGC target of 3e6 and a maximum ion injection time of 80 ms. There were 24 isolation windows for the two-part PulseDIA. The following MS2 analysis was performed at a resolution of 30,000, with an AGC target of 1e6 and a maximum ion injection time of 50 ms.

PulseDIA technology, in conjunction with high-resolution tandem MS, could provide broad coverage of identified and quantified peptides and proteins. Using two parts improved the coverage of the proteome and reduced the need for extensive fractionation. At the same time, the high resolution and accuracy of the Orbitrap Q Exactive HF (X) instrument enabled confident identifications and quantitations of peptides and proteins of interest.

#### 3.5 Data processing

##### 3.5.1 Hybrid spectral library construction.

We generated the hybrid library using FragPipe[5] (version 17.1) using an alternative “DIA_SpecLib_Quant” workflow. The PulseDIA files were converted to pseudo-MS/MS spectra using DIA-Umpire and then searched using the conventional MSFragger (DDA mode). All data, including DDA and pseudo-MS/MS spectra from DIA (Q1 files), were processed together using MSFragger (v.3.4), MSBooster, Percolator, ProteinProphet (Philosopher; v.4.2.1), and EasyPQP.

We selected a total of 5,503 files with high quality, including those generated with DDA and DIA, for the establishment of the spectral library. The main parameters of DIA-Umpire were set as follows: MS1 ppm: 10; MS2 ppm: 20; max Missed Scans: 1; we checked “Remove Background” and unchecked “Mass Defect Filter.” The FASTA file was downloaded from UniProt, a reviewed human Swiss-Prot sequence database (canonical and isoform: 51,548 entries). The decoys and contaminants were supplemented by Fragpipe, and the protein entry containing the ciRT peptide sequences was from Dropbox. For the library generation, “ciRT” was used for retention time correction using the default options. The library was filtered to a 1% false discovery rate (FDR) at the protein and peptide levels. The other parameters were set to their default values.

As shown in Figure 2d, the hybrid library contained 25,618,003 Protein Spectrum Matches (PSMs), 8,150,745 transitions, 196,892 peptides, and 12,768 protein groups (including 3,708 protein isoforms). The properties of the spectral library were characterized by DIALib-QC[6]. The precursor mass range covered 350–1250 m/z, most of the precursors were between 500 and 700 m/z, and the precursors primarily displayed two (48.2%) or three (38.4%) charges. The retention time correlation between two and three charges “ciRT” was high. Most peptides ranged between 8 and 20 amino acids, with a median of 11. The most common modification was methionine oxidation, detected in 42,545 peptides. Additionally, we detected 19,799 carbamidomethyled peptides at cysteine residues and 4,004 N-terminal acetylated. Almost 87.7% of proteins were detected with at least two proteotypic peptides, and 54.4% had more than eight proteotypic peptides. Also, 98.6% of peptides possessed over six fragment ions. Fragments from y-ions (61.3%) were more frequently detected than those from b-ions (38.7%). Finally, the fragment ion charge was mostly one (69.45%) or two (25.95%).

##### 3.5.2 DIA data analysis software and parameters

A total of 5648 DIA injections were analyzed by DIA-NN 1.8 using the hybrid MS spectral library we generated. The files included 4,766 brain samples, 308 pool samples, and 574 technical replicates. The parameters were set as follows: protease: Trypsin/P; missed cleavages: 1; N-term Methionine excision and Cysteine carbamidomethylation were set as variable modifications. The quantification was filtered to 1% FDR at the protein and peptide levels. The other parameters were left to their default values. Finally, we identified and quantified 149,777 peptides and 11,746 proteins (including 3,284 protein isoforms).

Proteomic data pre-processing is essential when analyzing proteomic data obtained from MS experiments. Here, the first step was to delete the samples overall with a low TIC signal or the protein identification quantity is lower than the median value of 80% of the overall identification quantity samples. Then proteins with over 80% missing values in all brain regions were considered unreliable and were removed from the dataset. The next step was to fill the remaining missing values. Considering that most missing values occurred when the protein content was below the detection threshold, imputation was performed by filling in all the missing values with [Dmin/10], where Dmin was the minimum of all available feature values in the proteome dataset, and one-tenth of Dmin was used to fill the missing values. This step ensured that all missing values were accounted for and allowed for consistent downstream analyses. Then quantile normalization[7] was employed to ensure that the distribution of protein intensities is similar across different samples, which reducing the impact of outliers and enabling fair comparisons between samples. Besides, the quantile normalization make the quantiles of each sample's distribution to a reference distribution.

Batch effects are a common issue in proteomic data analysis, particularly when samples are processed at different others by different instruments. The Limma R package is a widely used tool for removing batch effects; it uses a linear model to adjust for the batch effect while accounting for other sources of variation[8]. Here, the factor of sample preparation batches, instruments and acquisition durations was used to remove the batch effect with Limma.

#### 4. Quality Control Pipeline

To monitor and ensure the quality of our data, we set several checkpoints throughout our pipeline for generating data for our brain proteomics atlas (Figure S1j). The check steps Q1-Q3 ensured the accuracy of the collected brain samples and that the samples had been labeled correctly with their brain regions. For the proteomic data quality control, we set Q5-Q11 to monitor each step during the sample preparation and data generation. All these quality controls (QC) steps determined whether the data could be publicly available.

The major QC checkpoints during the process are marked in the following workflow: the Q in the green diamond indicates a QC step of the subsequent passage. All the steps are summarized in the table below.

| Checkpoints | Content | Criteria |
|---|---|---|
| Q1 | For all the sample, the integrity of the sample collection was assessed by | Samples must be >95% intact with no tearing, crushing, or deformation. |

| | | |
|---|---|---|
| | evaluating the presence of incompleteness, morphological abnormalities, or lack of white matter (especially in the cerebral cortex) within the samples. | Gross morphology should match expected neuroanatomy. The cerebral cortex must exhibit a clearly defined grey-white matter junction. Samples failing to meet these standards will be excluded from further processing. |
| Q2 | For all the brain tissue sections, we evaluated the integrity of the sections. Each section was checked for adhesion to the glass slide, tissue integrity, wrinkles, and drying artifacts. | Each tissue section must be fully adhered to the glass slide with no detachment. Tissue integrity must be preserved. Minor wrinkles are acceptable if they do not distort regional morphology. No drying artifacts are permitted under any circumstance. |
| Q3 | For the four healthy postmortem autopsy brain, the MRI data were evaluated for contrast, signal consistency across the dataset, and signal-to-noise ratio. We also checked whether the angle of the photos comprehensively covered all the sampling points. | |
| Q4 | Proteomic sample preparation QC. We used the K562 cell line sample (50,000 cells) to monitor the pipeline for the proteomic sample preparation. For each batch, we added one K562 sample, which was mixed and divided before evaluating the reproducibility of the sample preparation. | For each batch, replicate aliquots of the K562 control must show Pearson correlation >0.80 and median coefficient of variation (CV) <10% for quantified protein abundance. |
| Q5 | For all the sample, we assessing the quality of the tissues after dewaxing. This included evaluating the protein concentration and the integrity of the sample. | Samples must be fully dewaxed, indicated by tissue transparency and absence of visible paraffin residues. Protein concentration, measured via BCA assay, must be ≥0.5 µg/µL to ensure sufficient yield for downstream proteomic analysis. |
| Q6 | During the optimization process of the sample preparation pipeline, we evaluated the efficiency of the protein reduction and alkylation using MS: the presence of free sulfhydryl groups in the peptides indicates incomplete reduction and alkylation. By comparing the number of identified peptides between reduced and non-reduced peptides, we can evaluate the efficiency of the reduction and alkylation steps. | For each batch, four representative samples are assessed. The median proportion of identified peptides containing unmodified cysteine (free sulfhydryl groups) must be <5%. Additionally, no individual sample may exceed 10% unmodified cysteine peptides, ensuring complete reduction and alkylation efficiency during sample preparation. |
| Q7 | During the optimization process of the sample preparation pipeline, we assessing the quality of the peptides during digestion. The step includes the evaluation of the digestion efficiency and the enzyme specificity. The specificity of the protease and the missed cleavages can be evaluated by | Digestion efficiency and protease specificity are evaluated by MS analysis of peptides. Each sample must have a missed cleavage rate < 30%, meaning at least 70% of identified peptides contain two or fewer missed cleavages, ensuring effective and specific proteolytic digestion. |

| | | |
|---|---|---|
| | analyzing the digestion products using MS. By comparing the number of peptides with missed cleavages and the number of identified peptides, we can determine the missed cleavage rate and monitor the digestion efficiency. | |
| Q8 | The contamination of the samples with salts or insoluble suspended particles can negatively impact the accuracy of the quantification results. For all the sample, we used a multichannel UV spectrophotometer to detect A280, A230, and A260 to monitor the presence of impurities within the peptide samples. | Peptide samples are evaluated by UV spectrophotometry. Acceptable samples must have A260/A280 ratio < 0.8, indicating minimal nucleic acid contamination, and A230/A280 ratio < 0.5, reflecting low levels of salts, chaotropes, or organic solvents that could interfere with quantification accuracy. |
| Q9 | Weekly quality control of mass spectrometer. Before analyzing the samples, the instrument used for the MS analysis was usually calibrated and tested to ensure it was functioning correctly. This included testing the instrument's mass accuracy, resolution, and sensitivity. | Weekly, inject two randomized replicates of a standardized mouse liver digest. Mass accuracy must have a median error ≤5 ppm for precursors and ≤10 ppm for fragments. Each replicate must identify ≥80% of the historical median protein count (at 1% FDR) based on the last 10 successful QC runs. |
| Q10 | For each batch, to ensure the reproducibility of the data, it is often recommended to analyze the same sample (a mixture of different brain samples named “pool sample”) at the beginning. The same practice was applied to the technical and biological replicates. | Replicate measurements of the pool across the batch must show Pearson correlation >0.80 and median CV <10% for quantified protein abundance. |
| Q11 | For protein expression data, various QC metrics were used to assess the quality. These included evaluating the number of identified peptides and proteins, the peptide sequence coverage, and the distribution of peptide mass errors. Besides, the proteins with a high missing rate will be removed from the overall protein expression matrix, and the samples with low overall protein detection levels will be recollected the data. | Each sample must identify a minimum of 3,000 proteins. Proteins missing in more than 80% of samples are excluded from the expression matrix. Technical replicates must demonstrate Pearson correlation >0.80 and median CV <10% for quantified protein abundance, ensuring data consistency and reliability. |

The protein quantification was performed using both technical and biological replicates, leading to Pearson correlation coefficients of 0.96 and 0.97 and variable coefficients of 0.1 and 0.2, respectively (**Supplementary Fig. 2i**). The QC of the sample analysis did not indicate any significant batch effects between samples (**Supplementary Fig. 2j**).

#### 5. Allen Brain Atlas data processing

The transcriptome data from the Allen Institute retained all the cortical regions that could be registered to the Brainnetome brain atlas and all the subcortical nuclei. At the same time, we only kept transcripts that matched the FASTA files we used. Finally, 2113 brain regions and 18,415 transcripts were used for subsequent bioinformatics analyses.

To facilitate the analysis of HBA transcriptome data from the Allen Brain Institute along with our proteome data, we performed a re-annotation of the transcriptome samples based on the Brainnetome parcellation. The re-annotation process closely followed the previous section, with the key difference being the availability of coordinates in the MRI image space for each sample in the HBA dataset. Consequently, we initially mapped the Brainnetome atlas to the T1 image of each individual brain in the HBA dataset using nonlinear registration. Subsequently, the corresponding brain region for each sample was determined based on its coordinates in the T1 image space. We conducted a quality check through visual inspections to identify any apparent misassignments of brain regions for the samples.

#### 6. Epilepsy Sample Analysis

##### 6.1 Patient demographics and clinical information

Brain tissue specimens were obtained from 49 patients with medically refractory epilepsy undergoing therapeutic neurosurgical resection at our institution. All procedures were performed as part of standard clinical care for intractable epilepsy. Written informed consent was obtained from all patients or their legal guardians prior to surgery, and the study protocol received approval from the institutional ethics committee. A total of 100 gray matter samples were collected from six anatomical regions: temporal lobe (n=38), hippocampus (n=22), amygdala (n=19), frontal lobe (n=10), parietal lobe (n=8), and occipital lobe (n=3).

##### 6.2 Surgical tissue collection procedures

During neurosurgical resection, epileptogenic tissue was identified and excised. Resected specimens were immediately placed in sterile containers and transported to the laboratory on ice. Upon arrival, brain tissue was sectioned into small fragments, snap-frozen in liquid nitrogen, and stored at -80°C. To maintain methodological consistency with healthy brain tissue protocols, epileptic samples were processed within one month post-resection through formalin fixation (10% neutral buffered formalin) and paraffin embedding, generating formalin-fixed paraffin-embedded (FFPE) specimens for downstream proteomic analysis.

##### 6.3 Sample processing for proteomics

Sample preparation followed identical protocols to those employed for healthy brain FFPE specimens, utilizing pressure cycling technology (PCT)-assisted extraction methods.

##### 6.4 Differential expression analysis

Differential protein expression analysis was conducted between epileptic and healthy control samples across six anatomically distinct brain regions. Sample sizes varied by region: temporal lobe (epilepsy n=38, controls n=195), hippocampus (epilepsy n=22, controls n=47), amygdala (epilepsy n=19, controls n=10), frontal lobe (epilepsy n=10, controls n=369), parietal lobe (epilepsy n=8, controls n=178), and occipital lobe (epilepsy n=3, controls n=118). Following quality control assessment, proteins exhibiting >80% missing values across samples were excluded from downstream analyses. Statistical comparisons between epileptic and control groups were performed using Wilcoxon signed-rank tests. Differentially expressed proteins were identified using stringent criteria: fold change > 3 or < 1/3 and Benjamini-Hochberg adjusted p-value < 0.05. To identify consistently dysregulated proteins across brain regions, cross-regional protein overlap was visualized using the UpSetR package. Proteins demonstrating dysregulation across all six brain regions underwent biological pathway enrichment analysis using Ingenuity Pathway Analysis (IPA), with the top 10 significantly enriched pathways reported. Therapeutic target potential was assessed by cross-referencing dysregulated proteins with established drug target databases, including ChEMBL and DrugBank (accessed September 14, 2020).

#### 6.5 Assessment of age-related confounding factors in epilepsy proteomics.

A significant age disparity existed between healthy controls (n=4, aged 51-84 years) and epilepsy patients (n=49, aged 15-63 years). To evaluate age-related confounding effects, we conducted age-matched analysis using four temporal lobe epilepsy patients (aged 47-63 years) compared against four healthy temporal lobe samples. Statistical comparisons were performed using Wilcoxon signed-rank tests (**Supplementary Fig. 11a**). The age-matched cohort yielded 1541 differentially expressed proteins in temporal lobe tissue, defined by fold change > 3 or < 1/3 and adjusted p-value < 0.05 (**Supplementary Fig. 11b**). Comparative analysis revealed that age-stratified differentially expressed proteins represented a subset of those identified in the full patient cohort, with substantial overlap between age groups (**Supplementary Fig. 11c and d**). Cross-referencing with drug target databases identified 28 potential therapeutic targets, of which 22 were consistently detected across both age-matched (47-63 years) and full cohort (15-63 years) analyses in temporal lobe tissue. These findings demonstrate that age exerts limited confounding effects on epilepsy-associated protein dysregulation in the temporal lobe, validating the robustness of identified therapeutic targets for epilepsy treatment across different age populations.

#### 6.6 Network analysis (WGCNA)

**Network formation.** Weighted protein co-expression network analysis was performed using the WGCNA framework, following the pipeline for network construction and module identification. For the temporal lobe epilepsy (TLE) cohort (n = 38), proteins with more than 50% missing values were excluded, followed by filtering out proteins with low variability (median absolute deviation (MAD) < 0.35). One outlier sample was identified and removed via hierarchical clustering (method = "complete"). The final dataset comprised a 4635 × 37 $\log_2$-transformed protein abundance matrix, which had been corrected for covariates and batch effects. To determine an appropriate soft-thresholding power (β) for network construction, the pickSoftThreshold() function was employed over a power range of 2 to 20. A power of 7 was selected, based on the criterion that the resulting scale-free topology fit index (signed $R^2$) reached ≥ 0.8 and showed a plateauing trend, while preserving adequate mean connectivity. Network construction and module detection were performed using blockwiseModules() with the following parameters: power = 7, networkType = "signed", deepSplit = 2, minModuleSize = 60, mergeCutHeight = 0.15, TOMType = "signed", TOMDenom = "mean", minKMEtoStay = 0.30, reassignThreshold = 0.05, and corType = “bicor”. Pairwise biweight mid-correlations were computed between all protein pairs to generate a signed adjacency matrix, which was further transformed into a topological overlap matrix (TOM). The TOM encodes network interconnectedness by quantifying shared co-expression patterns, and 1-TOM was used as the dissimilarity metric for hierarchical clustering. Initial module delineation was conducted using dynamic tree cutting. Module membership (kME) was calculated as the Pearson correlation between each protein and the corresponding module eigengene. This process identified 10 modules, each containing ≥ 211 proteins. Modules with highly similar eigengenes (kME profiles) were merged using the mergeCloseModules() function (cutHeight = 0.4, corFnc = bicor), yielding 8 final modules. Following module merging, we recalculated signed kME values using the signedKME() function. To refine module assignments, proteins with intramodular kME < 0.28 were excluded, and unassigned ("gray") proteins with kME > 0.35 were reassigned to the module in which they exhibited the highest kME. This step accounted for the hybrid nature of WGCNA clustering, which is not solely based on correlation with eigengenes.

The same analytical pipeline was applied to the temporal lobe (TL) health control cohort (n = 195) and hippocampal epilepsy (HE) samples (n = 19). For both cohorts, proteins with >50% missingness, low MAD (<0.35), and outlier samples were removed. The TL health control network was constructed using 4604 proteins from 194 samples (power = 4, minimum module size = 50), while the HE network was based on 4887 proteins across 17 samples (power = 4, minimum module size = 20).

**Network preservation.** Module preservation across cohorts was evaluated using the WGCNA::modulePreservation() function, with the TLE network serving as the reference. Prior to analysis, proteins assigned to the “grey” module (i.e. unclustered) were excluded. Preservation statistics (Zsummary composite scores) were computed for the TLE network versus both TL health control and HE datasets, employing 500 permutations. To ensure reproducibility, a random seed of 1 was fixed and quickCor = 0 was specified; the analysis used a signed network with biweight mid-correlation (corFnc = "bicor"). Zsummary reflects combined preservation of module density and connectivity: values above 10 indicate strong preservation, values between 2 and 10 denote moderate evidence, and values below 2 suggest poor preservation.

**Module characterization.** Proteins exhibiting strong intramodular connectivity (kME > 0.6) within each TLE module were selected for pathway enrichment analysis. These high-kME proteins were analyzed using Ingenuity Pathway Analysis (IPA, Qiagen) to identify significantly overrepresented canonical pathways. We ranked the resulting pathways by p-value and focused on the top three most statistically significant. Among these, the pathway most closely linked to epilepsy pathophysiology, as supported by current literature, was selected to represent each module. Cell type–specific enrichment within each TLE protein module was evaluated using canonical marker sets from the Human Protein Atlas. We applied hypergeometric testing via the clusterProfiler::enricher() function, supplying custom TERM2GENE and TERM2NAME tables to assess over-representation of cell-type gene signatures. Module-specific protein lists were systematically compared against individual cell-type marker sets to identify significantly enriched associations across all defined cell categories.

#### 6.7 Potential drug targets for treating epilepsy.

To identify candidate drug targets in TLE, we applied a multi-step filtering strategy. First, differentially expressed proteins (epilepsy versus healthy control) exhibiting a fold change > 3 or < 1/3 with an adjusted p-value < 0.05 were filtered. From this subset, we retained proteins belonging to TLE-specific co-expression modules (module 4 or module 5), which are not preserved in TL healthy control networks. To ensure brain relevance, only proteins classified as brain-enriched by the Human Protein Atlas (https://www.proteinatlas.org/) were included. Finally, remaining proteins were cross-referenced against DrugBank (https://go.drugbank.com/) or ChEMBL(https://www.ebi.ac.uk/chembl/) to confirm druggability.

#### 7. Statistical and bioinformatic analyses

##### 7.1 Dimensionality reduction analysis

The analysis utilized a comprehensive proteomics dataset comprising 9771 proteins quantified across 2578 healthy control samples. Prior to dimensionality reduction, protein expression data underwent standardization and quality control filtering to ensure analytical robustness. Non-linear dimensionality reduction was performed using Uniform Manifold Approximation and Projection (UMAP) implemented through the uwot package in R. The analysis was conducted with the following optimized parameters: n_components = 2 for two-dimensional visualization, n_neighbors = 50 to balance local and global structure preservation, correlation distance metric to capture co-expression patterns, spread = 1 for uniform distribution of embedded points, and min_dist = 0.01 to allow tight clustering of similar samples. Z-score scaling was applied to standardize the input data matrix. Sample groupings were stratified by anatomical brain lobe regions to investigate regional proteomic signatures and potential clustering patterns. Linear dimensionality reduction was complemented by Principal Component Analysis using the proBatch::plot_PCA function. This analysis aimed to identify the primary sources of variance within the proteomic dataset and assess potential batch effects. Samples were grouped and color-coded according to individual MS instruments used for data acquisition to evaluate inter-instrument technical variance.

##### 7.2 Differential expression statistics

###### 7.2.1 Hemispheric and sex-specific regional proteomic analysis

Within each of the 121 defined gray-matter regions, we compared protein expression between hemispheres (left vs right) and between sexes (male vs female) within each hemisphere. Statistical comparisons were performed using Welch's two-sample t-tests, which do not assume equal variances between groups (t.test with paired = FALSE, var.equal = FALSE parameters). Analysis was restricted to samples derived from histologically normal gray matter across 121 discrete brain regions. Differential proteins were defined by a significance threshold of $p < 0.05$. For cerebellar regions, an additional criterion requiring a fold change > 3 or < 1/3 was applied to classify proteins as up- or down-regulated.

##### 7.3 Clustering algorithms (MOVICS)

###### 7.3.1 Data preprocessing and integration

The analysis included transcriptomic profiles covering 29,131 genes across 2141 gray matter samples and proteomic profiles quantifying 9771 proteins across 1522 gray matter samples. Sample metadata annotated each measurement to precise neuroanatomical locations using the Brainnetome atlas with hemispheric designations and donor identifiers. To mitigate technical artifacts, donor-specific batch effects in transcriptomic data were corrected via the removeBatchEffect algorithm (limma v3.54.1). Region-level expression signatures were derived by averaging measurements within each atlas-defined anatomical unit, yielding 214 non-cerebellar regions common to both omics layers. Feature selection prioritized molecular markers with established differential expression in cortical/subcortical structures, retaining 8129 RNA transcripts and 9473 proteins. Cerebellar regions were excluded to focus analysis on cortical/subcortical systems.

##### 7.3.2 Multi-omics integrative clustering

The optimal cluster number (k = 21) was determined by maximizing mean intra-cluster consensus scores computed via spectral clustering (k range: 2-30). Multi-omics clustering was performed using the MOVICS package (v0.99.17) with 10 algorithms: iClusterBayes, SNF, PINSPlus, NEMO, COCA, LRAcluster, ConsensusClustering, IntNMF, CIMLR, and MoCluster. All methods assumed Gaussian distributions for RNA/protein data and used default parameters. CIMLR was excluded from consensus due to non-convergence. A final consensus partition was derived by applying hierarchical clustering (euclidean distance, average linkage) to a consensus co-clustering matrix constructed from pairwise sample similarities across all algorithm-specific clustering solutions.

##### 7.3.3 Cluster validation and visualization

Cluster robustness was quantified via silhouette width analysis with function of getSilhouette. Anatomical associations were evaluated using Sankey diagram mapping cluster assignments to: Brainnetome atlas area-hemisphere combinations and lobular organization (frontal, temporal, parietal, limbic, occipital, insular, subcortical).

##### 7.3.4 Differential expression analysis

Transcriptomic and proteomic expression matrices underwent winsorization (3×IQR) to reduce outlier effects prior to differential analysis using the limma framework (runDEA function from MOVICS package). For each cluster versus all other clusters, features were classified as dysregulated markers if they satisfied three criteria: fold change > 3 or < 1/3, p-value < 0.05, and Benjamini-Hochberg adjusted p-value < 0.05.

#### 7.4 Cell type enrichment analysis

Cell type enrichment analysis was performed to delineate neurobiological underpinnings of molecular clusters, using canonical marker gene sets for 11 major brain cell types sourced from the Human Protein Atlas. Dysregulated signatures from each cluster-omics pair underwent over-representation analysis via Fisher's exact test implemented in the enricher function of clusterProfiler (v4.6.2).

#### 7.5 IPA pathway enrichment analysis

Canonical pathway enrichment was conducted on dysregulated signatures of each cluster using Ingenuity Pathway Analysis (IPA; QIAGEN). Significantly enriched pathways were defined by a threshold of p-value < 0.05, and conserved pathways were those observed in > 60% of clusters.

1 Zhu, T. *et al.* ProteomeExpert: a Docker image-based web server for exploring, modeling, visualizing and mining quantitative proteomic datasets. *Bioinformatics* **37**, 273-275, doi:10.1093/bioinformatics/btaa1088 (2021).
2 Cai, X. *et al.* High-throughput proteomic sample preparation using pressure cycling technology. *Nat Protoc* **17**, 2307-2325, doi:10.1038/s41596-022-00727-1 (2022).
3 Gao, H. *et al.* Accelerated Lysis and Proteolytic Digestion of Biopsy-Level Fresh-Frozen and FFPE Tissue Samples Using Pressure Cycling Technology. *J Proteome Res* **19**, 1982-1990, doi:10.1021/acs.jproteome.9b00790 (2020).

4 Cai, X. *et al.* PulseDIA: Data-Independent Acquisition Mass Spectrometry Using Multi-Injection Pulsed Gas-Phase Fractionation. *J Proteome Res* **20**, 279-288, doi:10.1021/acs.jproteome.0c00381 (2021).

5 Kong, A. T., Leprevost, F. V., Avtonomov, D. M., Mellacheruvu, D. & Nesvizhskii, A. I. MSFragger: ultrafast and comprehensive peptide identification in mass spectrometry-based proteomics. *Nat Methods* **14**, 513-520, doi:10.1038/nmeth.4256 (2017).

6 Midha, M. K. *et al.* DIALib-QC an assessment tool for spectral libraries in data-independent acquisition proteomics. *Nat Commun* **11**, 5251, doi:10.1038/s41467-020-18901-y (2020).

7 van Hijfte, L. *et al.* Alternative normalization and analysis pipeline to address systematic bias in NanoString GeoMx Digital Spatial Profiling data. *iScience* **26**, 105760, doi:10.1016/j.isci.2022.105760 (2023).

8 Ritchie, M. E. *et al.* limma powers differential expression analyses for RNA-sequencing and microarray studies. *Nucleic Acids Res* **43**, e47, doi:10.1093/nar/gkv007 (2015).

### 西湖大学伦理初始审查申请与审批表

编号：20240527 GTN002　　　　　　　　日期：2024 年 05 月 27 日

| |
|---|
| 项目名称：基于质谱的人脑空间蛋白质组研究 |
| 项目承担单位：　西湖实验室（生命科学和生物医学浙江省实验室）<br>项目负责人：郭天南 |
| 项目合作单位：上海交通大学基础医学院　　　合作联系人：李岩 |
| 伦理委员会审查资料：<br><br>■实验方案　　☐知情同意书　　■项目负责人信息<br>☐项目相关机构的合法资质证明（仅临床药物实验需提供）<br>☐其他资料：＿＿＿＿＿＿＿＿＿＿ |
| 涉及人或动物的研究内容和研究方案摘要（可另附具体实验方案）：<br>本研究计划使用目前国际领先的质谱仪器和基于 MRI 的功能脑分区图谱-Brainnetome Atlas7，对人脑组织进行系统的高空间分辨率的蛋白质组分析，建立一个详尽的人脑蛋白质组表达图谱资源库，系统比较生理和病理状态的脑组织的蛋白质表达差异。此外结合膨胀胶技术开发一种操作简单且可以完成完整细胞取材的单细胞取样方法 8。传统的蛋白质组研究需要大量的样本处理和实验操作，工作量巨大且容易出现误差。利用自动化的样本采集和前处理流程，可以大大提高研究效率，减少人为误差，确保数据的准确性和可靠性。 |

| |
|---|
| 研究需要使用的生物样本来源类型：<br>☐招募受试者采集生物样本 （需额外提供受试者知情同意书）<br>■采用以往保存的生物样本<br>☐采用实验动物的生物样本<br>☐其他来源：________________________________ |
| 研究需要使用生物样本的具体来源及计划处置方式：<br>来自遗体捐献中心的福尔马林固定的健康人脑样本进行 MRI 数据采集，对全脑进行功能分区。使用穿刺针对全脑进行采样。样本采集后的人脑重新进行 MRI 扫描确定所采集的样本空间位置坐标并对所有样本进行区域标注。标注后的样本进行蛋白质组学数据采集和分析。 |
| 项目负责人承诺：<br>本人将投入足够的时间和精力开展本项目研究，确保项目的研究进度和质量，并将遵循实验方案及伦理委员会的要求。<br>项目负责人：郭天南<br>2024年 5月 27日 |

伦理委员会审查意见：

☑批准　　☐不批准　　☐修改后再审

☐暂停研究　　☐终止研究

审查意见详述：

有效期3年.

西湖大学伦理委员会（盖章）

主任签字：杨剑 SLS

2024 年 6 月 7 日

### 涉及人体科研项目伦理委员会批准函

### RESEARCH EHTICS COMMITTEE APPROVAL FORM

| 项目名称 Project Name | 人脑组织资源库协作网络平台 | | | 项目受理号 Project Pending No. | （2022）伦审第（A-01）号 |
|---|---|---|---|---|---|
| 项目类别 Project Category | ☑基础 Basic ☐临床 Clinical ☐其他 Other | | | 样本例数 Sample Size | 50 |
| 项目来源 Project Source | 科技部 | | | 研究期限 Study Duration | 2021.09-2026.12 |
| 项目负责人 Principal Investigator | 徐天乐 | 研究系室 Research Department | 解剖学与生理学系 | 职 称 Title | 教授 |

**伦理审查方式：** ☐会议审查 Full Board Review ☑快速审查 Expedited Review ☐紧急会议审查 Emergency Review

**持续审查及其频率：** ☑是 Yes： ☐ 3 个月 Three months ☐ 6 个月 Six months ☑ 1 年 One year ☐否 No

**下面划☑的研究相关文件已经审阅**
The following documents have been reviewed and approved by the ethics committee：

☑伦理审查申请报告 Application Report of ethical review
☑课题研究方案 Research proposal
☑知情同意书 Informed Consent Form
☑样本来源证明 Proof of sample source
☑研究者简历 PI's CV
☐其他 Others

1、**审阅结果** Review Result

☑同意 Approve

2、**审批意见** Review Comments

**该项目经上海交通大学基础医学院科研伦理委员会审核，其研究内容及方案设计基本符合伦理规范，同意按申报方案进行研究。**

邵强

主任委员（签名）
Signature of Chairman

上海交通大学基础医学院伦理委员会（盖章）
（印章：上海交通大学基础医学院 科研伦理委员会）

日期（2022 年 3 月 31 日）
Date (YYYY / MM/ DD)

地址：上海市重庆南路 227 号
邮编：200025
电话：021-63846590-776169

Address: No. 227 South Chongqing Road, Shanghai, P.R China
Post Code: 200025

### 伦理审查同意函

#### HIRB Approval Letter

**声明**：本伦理审查委员会按照中国 GCP 和有关法规组成和工作，其审查和工作过程不受伦理审查委员会以外任何组织及个人的影响

**批件号：（2022）临审第（590）号修正 2**

| | |
|---|---|
| 审查会议日期 | 2022 年 10 月 25 日 |
| 审查会议地点 | 腾讯视频会议 |
| 研究项目名称 | 人脑皮层结构的多组学研究 |
| 审查文件 | 1. 修正案申请报告<br>2. 研究修改处总结<br>3. 研究方案（03，2022 年 7 月 29 日）<br>4. 知情同意书（02，2022 年 7 月 29 日） |
| 申办者 | 无 |
| 临床研究单位 | 复旦大学附属华山医院神经外科 |
| 主要研究者 | 陈 亮 |
| 伦理审查方式 | ■ 会议审查　口快速审查　口紧急会议审查 |
| 审查委员 | 伍蓉（研究员，女），顾勇（医学教授），倪泉兴（医学教授），冯晓源（医学教授），王大猷（主任药师），董强（医学教授），杨勤萍（医学教授、女）、王坚（医学教授），丁玎（副研究员、女），曹国英（副主任药师，女），陈建波（律师，外单位），王跃娟（非医药专业，女，外单位），刘海涛（伦理学，女，外单位），钟良（医学教授），黄海辉（医学教授，女），吴翠云（助理研究员，女） |
| 审查意见 | 1. 经本伦理审查委员会审查：**同意按修正案进行研究。**<br>**意见和建议：无**<br>2. 伦理审查委员会对该研究实施过程的年度/定期审查：■是　口否<br>审查频度为研究首次同意之日起：口 3 个月　口 6 个月　■1 年<br>3. 伦理审查委员会有权根据实际进展情况改变年度/定期审查频度。 |

主任委员或副主任委员签字：[signature]

复旦大学附属华山医院伦理审查委员会（盖章）

日期：2022.11.2

**注意：（请仔细阅读）**

1. 凡是涉及人类遗传资源保护或者按照国家规定必须经有关部门专项审批的内容，均必须在项目执行前向有关部门申报并获得同意。
2. 本伦理审查委员会同意的项目均为涉及人的生物医学研究，必须严格按照所同意方案规定的期限和受试者例数完成，不得随意超过。如需作为临床项目常规开展，必须经医院医学伦理委员会同意，并按照国家新技术申报相关规定向有关部门申请同意。
3. 本同意函可能在各中心机构及其伦理委员会备案。如果对方案在贵机构的可行性（包括研究者的资格与经验、设备与条件等）有不同意见，请及时与本伦理审查委员会联系。
4. 在第 1 例受试者入组之前，研究信息应在公众所及的网站上登记，如我国医学研究登记备案信息系统。
5. 已同意项目须遵循本伦理审查委员会同意的方案执行，须符合国家各部委、NMPA 相关法规指南和《赫尔辛基宣言》等我国认可的国际指南规定的伦理原则。
6. 暂停/提前终止临床研究，请及时通知伦理审查委员会。
7. 发生可疑且非预期的严重不良反应，须经研究者快速报告本伦理审查委员会。
8. 对已获得伦理审查同意的临床研究方案、知情同意书等材料的修改及主要研究者更换等，须及时递交本伦理审查委员会，获得审查同意后方可执行，但为了及时消除对受试者的紧急危害（事后及时报告伦理审查委员会）或更换监查员、电话号码等仅涉及临床试验管理方面的改动除外。
9. 发生增加受试者风险或显著影响研究实施的不依从/偏离方案事件，或为了避免对受试者的即刻危险偏离方案，应及时报告本伦理审查委员会；其他一般不依从/偏离方案事件可定期报告。
10. 根据伦理审查委员会对年度/定期审查频率的意见，无论研究开始与否，请在年度/定期审查日到期前 1 个月提出审查申请。
11. 完成临床研究，须提交结题报告给本伦理审查委员会。

地址：上海市乌鲁木齐中路 12 号；邮编：200040；电话：021-52888045

### 人脑皮层结构的多组学研究

**主要研究者：陈亮**

**临床研究科室：**复旦大学附属华山医院神经外科

**申办者：**无

**方案号：KY2022-590**

**方案版本号：**03，2022 年 7 月 29 日

主要研究者签名：

日期：2022 年 5 月 7 日

### 主要研究者联系方式

**复旦大学附属华山医院神经外科**


姓名：陈亮

职称：主任医师

地址：上海市静安区乌鲁木齐中路 12 号复旦大学附属华山医院神经外科

| | |
|---|---|
| 电话号码（办公室） | **021-52888771** |
| 手机 | **13311735889** |
| 电子邮箱 | |


### 研究小组成员联系方式

**复旦大学附属华山医院神经外科**


姓名：毛颖

职称：主任医师

地址：上海市静安区乌鲁木齐中路 12 号复旦大学附属华山医院神经外科

| | |
|---|---|
| 电话号码（办公室） | **021-52888771** |
| 手机 | **13801769152** |
| 电子邮箱 | |

姓名：杨辉

职称：青年研究员

地址：上海市静安区乌鲁木齐中路 12 号复旦大学附属华山医院神经外科

| | |
|---|---|
| 电话号码（办公室） | **021-54237056** |
| 手机 | **18626215485** |
| 电子邮箱 | |


**中科院脑科学与智能技术卓越创新中心**


姓名：李澄宇

职称：研究员

地址：上海市岳阳路 320 号中科院脑科学与智能技术卓越创新中心生命科学实验楼

| | |
|---|---|
| 电话号码（办公室） | **021-54921824** |
| 手机 | **15021632759** |
| 电子邮箱 | |

**复旦大学脑科学转化研究院**

姓名：舒友生

职称：研究员

地址：上海市徐汇区东安路 130 号

| | |
|---|---|
| 电话号码（办公室） | **021-54237723** |
| 手机 | **13810372549** |
| 电子邮箱 | |


**复旦大学脑科学研究院**

姓名：诸颖

职称：青年研究员

地址：上海市徐汇区东安路 130 号

| | |
|---|---|
| 电话号码（办公室） | **021-54237688** |
| 手机 | **13395750276** |
| 电子邮箱 | |

姓名：侯伟贞

职称：博士研究生

地址：上海市徐汇区东安路 130 号

| | |
|---|---|
| 电话号码（办公室） | **021-54237688** |
| 手机 | **19121792676** |
| 电子邮箱 | |

姓名：何炜

职称：博士研究生

地址：上海市徐汇区东安路 130 号

| | |
|---|---|
| 电话号码（办公室） | **021-54237688** |
| 手机 | **13725395208** |
| 电子邮箱 | |

姓名：赵曼曼

职称：博士研究生

地址：上海市徐汇区东安路 130 号

| | |
|---|---|
| 电话号码（办公室） | **19121803482** |
| 电子邮箱 | **** |


**复旦大学附属华山医院神经外科**

姓名：胡杰

职称：主任医师

地址：上海市静安区乌鲁木齐中路 12 号复旦大学附属华山医院神经外科

| | |
|---|---|
| 电话号码（办公室） | **021-96567576** |

| | |
|---|---|
| 电子邮箱 | |


姓名：杨伯捷

职称：主任医师

地址：上海市静安区乌鲁木齐中路 12 号复旦大学附属华山医院神经外科

| | |
|---|---|
| 手机 | **13901613855** |
| 电子邮箱 | **** |

姓名：赵帆

职称：副主任医师

地址：上海市静安区乌鲁木齐中路 12 号复旦大学附属华山医院神经外科

| | |
|---|---|
| 电话号码（办公室） | **021-52888771** |
| 手机 | **13901778205** |
| 电子邮箱 | |

姓名：史之峰

职称：副主任医师

地址：上海市静安区乌鲁木齐中路 12 号复旦大学附属华山医院神经外科

| | |
|---|---|
| 手机 | **13917793493** |
| 电子邮箱 | **** |

姓名：邹翔

职称：主治医师

地址：上海市静安区乌鲁木齐中路 12 号复旦大学附属华山医院神经外科

| | |
|---|---|
| 电话号码（办公室） | **021-52888771** |
| 手机 | **13611633383** |
| 电子邮箱 | |

姓名：岳琪

职称：主治医师

地址：上海市静安区乌鲁木齐中路 12 号复旦大学附属华山医院神经外科

| | |
|---|---|
| 电话号码（办公室） | **021-52888771** |
| 手机 | **13916162867** |
| 电子邮箱 | |

姓名：齐曾鑫

职称：主治医师

地址：上海市静安区乌鲁木齐中路 12 号复旦大学附属华山医院神经外科邮编：**200040**

| | |
|---|---|
| 电话号码（办公室） | **021-52887030** |
| 手机 | **17317709785** |
| 电子邮箱 | |

| | |
|---|---|
| 姓名：沈超 | |
| 职称：主治医师 | |
| 地址：上海市静安区乌鲁木齐中路 12 号复旦大学附属华山医院神经外科邮编：**200040** | |
| 电话号码（办公室） | **021-52887030** |
| 手机 | **13917182144** |
| 电子邮箱 | **** |

**目录**

**研究小组成员** **2-5**

**临床研究方案** **7-13**

**参考文献** **14-15**

## 正文

##### 1. 研究背景

“什么使我们成为人类”一直是吸引人类探索自身的重要科学问题。细胞是生命的基本单元。其中，作为人类现知最复杂的器官——人类大脑，大约由上千亿个各种类型的细胞组成。然而，迄今为止，我们对于人类大脑中的细胞类型，及其分子标识，以及它们在脑中的空间定位仍知之甚少。绘制人类大脑细胞类型和空间转录组图谱将有助于我们解析大脑中各种细胞类型和由此形成的复杂网络的功能，极大地促进我们从分子和细胞层面探究人类大脑的奥秘。随着中国老龄人口的增加，中国社会正在快速进入老龄化社会。老年人口的增多，极大的增加了我们对于理解衰老如何发生的需求，只有真正理解衰老的发生过程及机制，才能很好的预防治疗各种衰老相关的疾病。在众多衰老影响的疾病中，阿尔兹海默症是几个重点待研究疾病之一。阿尔兹海默症病影响了众多老年人的身体健康，并且发病机制未知。我们尝试从衰老的角度去探究阿尔兹海默症的发病机制。

人类和非人灵长类动物在认知和行为能力上存在着明显的差异，如句法语法语言、符号思维、自我反省、长期规划能力、自传体记忆、心理理论和艺术创作能力等都是人类特有的能力和行为[1-3]。大脑作为人类身份的核心组成部分，理解其结构和功能对于探索人类异性的行为和能力至关重要。人类大脑的非凡能力是区分人类与其他类人猿的最明显的特征之一。然而，我们对神经系统，特别是大脑在人类进化谱系中如何变化的理解仍然非常有限。

对人类和非人灵长类动物大脑的比较研究很难以传统的研究方法实现。由于伦理和法律的限制，实验生物学中使用的大多数方法不适用于研究人类和非人灵长类大脑。因此，我们对控制灵长类大脑发育和功能的分子和细胞机制的大部分理解，都来源于对一些遗传距离较远的模式生物的实验研究。虽然这些模型系统的使用使生物医学科学发生了革命性的变化，但是从它们中推断出来的结论有时候并不适用于类人猿(4-6)。

近年来，测序技术的发展，为研究人脑和非人灵长类动物大脑开启了新的大门。例如通过物种间全基因组比较，发现了人类加速进化的基因和调控序列，且神经系统基因的进化突变富集于调控序列，说明人类和非人灵长类之间的许多表型差异可能来源于调控序列的变异(7)。而调控序列的变异往往通过影响基因表达来发挥作用。因此近一步的一系列研究通过脑组织的转录组分析，寻找在人脑中特异性表达的基因及其功能(8-14)。申请人前期的工作通过比较人、黑猩猩和恒河猴大脑多脑区的转录组，发现了成年人脑在分子和细胞水平上的重构(13)，及其发育基础(14)。

本临床研究项目拟通过对成年人脑的多个脑区进行多组学测序（单细胞组学，蛋白质组、结合膜片钳电生理的 Patch-seq 等），并与非人灵长类进行比较，探索人脑特异性基因表达的调控机制，特别解析非编码区序列变化对基因表达的影响。

**2. 研究目的**

本实验希望通过对成年人脑的多个脑区运用单细胞测序、时空芯片测序、多重荧光原位杂交、电生理、Patch-seq 以及生物信息分析等前沿技术，对人类大脑中多个脑区中的细胞类型和联接、基因表达谱及其空间定位进行普查式分析，发现人类大脑所具有独特的细胞类型和突触联接，绘制出人类大脑细胞类型和空间转录图谱；并与非人灵长类进行比较，探索人脑特异性基因表达的的特点及其调控机制，特别解析非编码区序列变化对基因表达的影响；运用该图谱揭示人类大脑中细胞类型的组织结构关系，特别是不同细胞类型与其邻近细胞的关系；为人类大脑内的细胞构成和空间关系提供一个详实的参考数据库，促进脑疾病的诊疗，为脑健康做出积极的贡献。

**3. 研究设计**

本研究为基础研究实验，拟初步招募大脑或小脑皮层有癫痫、出血或肿瘤性病灶需要切除的病人 400 名。本实验无随机化分组设置，不设盲，无干预因素。

**4. 受试者的入选标准，排除标准，中止标准**

**入选标准**：受试者必须符合以下所有标准：

1) 因脑出血、脑外伤、脑肿瘤、癫痫需要手术的患者，且术前判断手术标本有可能累及大脑或小脑皮层的患者。
2) 脑 MRI 无其他严重神经系统疾病证据。
3) 经研究者确认无重大慢性疾病、其他恶性肿瘤或急性传染病等。
4) 须由受试者或其法定监护人书面签署知情同意书。
5) 受试者无特殊年龄或性别限定

**排除标准**：符合以下任何一项标准的受试者将被排除在研究之外：

1) 临床报告有其他严重神经或精神疾病或胃肠道、心血管、肝脏、肾脏、血液系统、肿瘤内分泌、呼吸系统、免疫缺陷及其他严重疾病。药物滥用迹象或任何长时间痛苦状况（昏迷、长时间发热、缺氧、长时间脱水、低血糖、和多器官衰竭）。

2) 具有染色体或大规模基因组异常。

3) 脑组织有明显的畸形，包括但不限于神经管、前脑、脑干、颅神经、小脑、脊髓异常等。

4) HIV、乙肝（Hepatitis B）、丙肝（Hepatitis C）等病毒感染未愈者。

**中止标准：**符合以下任何一项标准的受试者将被立刻中止研究：

1) 受试者不愿或不能继续参加试验者；

2) 发生了不可耐受的不良事件或严重不良事件，研究者判断继续参加试验对受试者经受的风险大于其获益者；

3) 误纳入的受试者；

4) 试验过程中严重违背试验方案；

5) 无任何可评价记录（临床发作次数、脑电图检查等）。

**5. 根据统计学原理计算要达到研究预期目的所需的病例数**

目前研究对于不同脑区，在保证脑片活性或样本生物活性的前提下，Patch-seq 实验部分中，重复记录 5 个以上不同样本脑片的电生理特性；免疫组化部分和组学研究部分，需 5 个以上不同样本脑片，需患者 400 例。

**6. 研究流程**

1）选择受试者，了解患者基本病史，服用药物情况，检查结果，进行临床评估，告知受试者注意事项并签署知情同意书。本研究需要收集大脑或小脑皮层有癫痫、出血或肿瘤性病灶需要切除的病人 400 名。

2）根据患者临床诊治需要按照临床诊治流程，行手术切除。

3）在不影响患者疾病诊治的前提下，从手术切除样本中进行采样，每块手术切除样本采集标本大小约 0.5-1$cm^3$，用提前配备好的标本保存液转移采集标本至实验室。

4）将采集样本进行切片孵育，离体电生理记录，记录样本电生理特性。

5）将采集样本进行包括单细胞转录组，空间转录组，表观遗传组，蛋白质组在内的多组学测序。

6）整合分析数据

试验流程图：

选择受试者，了解病史、服药情况和既往检查，进行临床评估，告知受试者注意事项，签署知情同意书；

↓

手术切除

↓

从手术标本采样，用提前配备好标本保存液转移手术切除标本至实验室

↓

实验室切片、孵育

↓

进行样本离体电生理记录

↓

进行多组学测序

↓

数据整合分析

**7. 中止研究的规定：**

结束标准：出现以下任何一项，研究者将立即停止所有研究

a. 研究方案设计失误

b. 研究风险较大，伦理委员会审查后停止该试验

c. 研究无价值

**8. 受试者的编码、随机数字表及病例报告表的保存手续；**

受试者按照入组顺序及疾病类型进行编号(如 EP-01,EP-02)。研究记录表由专人填写，电子版保存在研究团队数据分析电脑。 所有资料保留至研究结束 10 年。

**9.安全性事件的管理**

本研究仅收集相关手术患者的手术切除标本，不涉及相应的不良事件。

**10. 统计分析计划，统计分析数据集的定义和选择；**

所有统计分析均采用 SPSS 22.0 统计分析软件进行。

**11. 数据管理、数据保密和数据可溯源性的规定；**

（1）研究记录表的构建

数据管理员根据研究方案构建研究记录表

（2）数据的录入与修改

数据的录入与修改由研究者完成，数据应来源与原始记录和实验室检查报告单等原始文件并英语原始文件一致，试验中的任何观察、检查结果均应及时、正确、完整、清晰、规范、真实的录入表格或数据库中。

（3）研究参与者（受试者）信息保密计划

所有研究参与者（受试者）的信息必须严格保密，参与研究及在研究中的个人资料均属保密范围。研究参与者（受试者）的信息和研究数据将以研究编号数字而非其名字加以表识。可以识别其身份的信息将不会透露给研究小组以外的成员，除非获得研究参与者（受试者）的许可。所有的研究成员都被要求对研究参与者（受试者）的身份保密。研究参与者（受试者）的档案将保存在有锁的档案柜中，仅供研究人员查阅。为确保研究按照规定进行，必要时，政府管理部门或者伦理审查委员会的成员按规定可以在研究单位查阅研究参与者（受试者）的个人资料。这项研究结果发表时，将不会披露研究参与者（受试者）个人的任何资料。华山医院提供给复旦大学脑科学研究院的样本和数据必须进行编码和去标识处理。

（4）研究数据的保密计划

研究数据同属保密范围，所有研究成员都将被要求对研究数据保密，不得未经主要研究者许可将研究数据告知课题组以外的成员，华山医院提供给复旦大学脑科学研究院的样本和数据，复旦大学脑科学研究院使用完毕之后应当返还，同时未经医院许可不得再做其他用途，和提供给其他人或单位。

**12. 临床研究的质量控制与质量保证；**

临床试验的质量控制系指用以保证与临床试验相关活动的质量达到要求的操作性技术和规程。主要通过制定临床试验标准操作规程（standard operation procedure，SOP），确保临床试验自始至终遵循 SOP 的操作规程。

临床试验前应对所有参试人员进行相关的 SOP 培训，并在试验开始阶段认真监察 SOP 的执行，在执行中应对 SOP 的适用性和有效性进行系统的检查，对确认不适用的 SOP 进行修改和补充。

主要研究者组织相关培训，确保研究规范进行，研究记录表的填写和其它报告都遵循

GCP 原则和研究方案。所有数据和资料必须可以被查证；为了保证研究数据的可靠性，所有观察和发现应具有可验证性。研究的每一阶段都将运用质量控制来保证所有数据的可靠和研究程序的准确。主要研究者应确保研究者遵守试验方案，确认数据的准确及记录报告的完整，确保在研究开始前取得所有受试者的知情同意书。任何违背或偏离方案应及时报告伦理委员会。

**13. 研究相关的伦理学；**

本临床试验应当遵守《赫尔辛基宣言》及相应的法律法规。对每位捐赠术手术切除组织的患者应明确告知其组织将用于本项目研究。该研究将在获得复旦大学附属华山医院伦理委员会的批准后开展。

**14. 受试者招募方法和获得知情同意书过程；**

对受试者招募通过具有经验的神经外科专科医师在神经外科门诊招募。 知情同意书的签署遵循以下原则及事项：1. 必须向受试者提供有关试验的详细情况（包括试验目的、受试者预期的受益与承担的风险），并获得受试者同意。 2. 受试者参加试验是自愿的，且在试验的任何阶段均有权随时退出试验而不会遭到歧视或报复。3. 必须给受试者充分的考虑时间以便决定是否愿意参加。4. 受试者在充分了解试验详细情况的基础上， 愿意参加试验，由受试者或其合法代表在同意书上签字并注明日期。5. 受试者在试验过程中的个人资料严格保密。 6. 必须向受试者说明发生试验相关的损害时可以获得适当的治疗或补偿。7. 试验期间研究者必须随时了解受试者有关的信息资料。 8. 如有新的重要的有关试验的资料可能涉及受试者的同意，则必须将知情同意书作书面修改，送伦理委员会批准后，征得受试者同意。

**15. 研究预期的进度和完成日期；**

本临床研究项目预计执行期限 5 年。

（1）2022 年 6 月至 2027 年 4 月完成所有脑区样本募集及电生理实验、多组学测序。完成切片、染色等实验

（2）2024 年 5 月至 2027 年 4 月 汇总实验数据，完成数据分析，完成研究论文报告，初步研究成果，总结报告，文章撰稿。

**16. 研究结束后的随访和医疗措施；**

本研究仅需手术病人的手术切除组织标本与相应的病理结果，不涉及研究结束的随访和医疗措施。

**17. 各方承担的职责及其他有关规定；**

本研究项目将采用的受试者生物样本来源于手术切中的病灶或灶旁组织。由复旦大学华山医院外科医生对受试者病灶和灶旁组织进行取材。在使用手术切除的病灶或灶旁组织时，捐献者或其近亲属已签署知情同意书，同意将样本用于各项科学研究。华山医院提供给复旦大学脑科学研究院、复旦大学脑科学转化研究院、中科院脑科学与智能技术卓越创新中心的样本和数据必须进行编码和去标识处理。华山医院提供的样本和数据，仅限于本课题使用，使用完毕之后应当返还，同时未经医院许可不得再做其他用途，和提供给其他人或单位。

华山医院负责对项目进行统筹规划，招募受试者，进行受试者病灶和灶旁组织进行取材，同时对取得的样本进行多组学测序及数据分析，免疫组化验证。

复旦大学脑科学研究院、复旦大学脑科学转化研究院、中科院脑科学与智能技术卓越创新中心负责对取得的组织样本进行离体电生理检测，多组学检测，协助完成测序数据分析及免疫组化验证。

**22. 参考文献**

1. P. Lieberman, The evolution of language and thought. J Anthropol Sci 94, 127-146 (2016).
2. R. E. Passingham, What is special about the human brain? , Oxford psychology series (Oxford University Press, Oxford ; New York, 2008), pp. xii, 269 p.
3. D. C. Penn, K. J. Holyoak, D. J. Povinelli, Darwin's triumph: Explaining the uniqueness of the human mind without a deus ex machina. Behav Brain Sci 31, 153-178 (2008).
4. Z. Petanjek et al., Extraordinary neoteny of synaptic spines in the human prefrontal cortex. P Natl Acad Sci USA 108, 13281-13286 (2011).
5. B. I. Bae, D. Jayaraman, C. A. Walsh, Genetic Changes Shaping the Human Brain. Dev Cell 32, 423-434 (2015).
6. K. Semendeferi et al., Spatial organization of neurons in the frontal pole sets humans apart from great apes. Cereb Cortex 21, 1485-1497 (2011).
7. R. Haygood, C. C. Babbitt, O. Fedrigo, G. A. Wray, Contrasts between adaptive coding and noncoding changes during human evolution. P Natl Acad Sci USA 107, 7853-7857 (2010).
8. P. Khaitovich et al., Regional patterns of gene expression in human and chimpanzee brains. Genome Res 14, 1462-1473 (2004).
9. M. C. Oldham, S. Horvath, D. H. Geschwind, Conservation and evolution of gene coexpression networks in human and chimpanzee brains. Proc Natl Acad Sci U S A 103, 17973-17978 (2006).
10. M. Somel, X. L. Liu, P. Khaitovich, Human brain evolution: transcripts, metabolites and their regulators. Nat Rev Neurosci 14, 112-127 (2013).
11. X. Liu et al., Extension of cortical synaptic development distinguishes humans from chimpanzees and macaques. Genome Res 22, 611-622 (2012).
12. Z. S. He, H. Bammann, D. D. Han, G. C. Xie, P. Khaitovich, Conserved expression of lincRNA during human and macaque prefrontal cortex development and maturation. Rna 20, 1103-1111 (2014).
13. A. M. M. Sousa et al., Molecular and cellular reorganization of neural circuits in the human lineage. Science 358, 1027-1032 (2017).
14. Y. Zhu et al., Spatiotemporal transcriptomic divergence across human and macaque brain development. Science 362, 1267-+ (2018).
15. Y. Zhu, M. Li, A. M. Sousa, N. Sestan, XSAnno: a framework for building ortholog models in cross-species transcriptome comparisons. BMC Genomics 15, 343 (2014).

附表 1

**1.Zhu Y#, Sousa AMM#, Gao T#, Skarica M#, Li M#, Santpere G, Esteller-Cucala P, Juan D, Ferrández-Peral L, Gulden FO, Yang M, Miller DJ, Marques-Bonet T, Kawasawa YI, Zhao H, Sestan N* (2018). Spatio-temporal transcriptomic divergence across human and macaque brain development. Science. 362(6420):eaat8077. (*Co-first author)**

**2.Sousa AMM#, Zhu Y#, Raghanti MA, Kitchen RR, Onorati M, Tebbenkamp ATN, Stutz B, Meyer KA, Li M, Kawasawa YI, Liu F, Perez RG, Mele M, Carvalho T, Skarica M, Gulden FO, Pletikos M, Shibata A, Stephenson AR, Edles MK, Ely JJ, Elsworth JD, Horvath TL, Hof PR, Mane SM, Noonan JP, State MW, Lein ES, Knowles JA, Marques-Bonet T, Sherwood CC, Gerstein MB, Sestan N* (2017). Molecular and cellular reorganization of neural circuits in the human lineage. Science. 358, 1027-1032. (*Co-first author)**

**3.Zhu Y#, Silbereis JC#, Pochareddy S#, Li M, Sestan N*. (2016) The Molecular Landscape of the Developing Human Central Nervous System. In Genomics, circuits, and pathways in clinical neuropsychiatry, T. Lehner, B. Miller, and M. State, eds. (Boston, MA: Elsevier). pp. 203-220. (*Co-first author； Book Chapter)**

**4.Zhu Y, Li M, Sousa AM, Sestan N*(2014). XSAnno: a framework for building ortholog models in cross-species transcriptome comparisons. BMC Genomics. 15, 343.**

**5.Kang HJ#, Kawasawa YI#, Cheng F#, Zhu Y#, Xu X#, Li M#, Sousa AM, Pletikos M, Meyer KA, Sedmak G, Guennel T, Shin Y, Johnson MB, Krsnik Z, Mayer S, Fertuzinhos S, Umlauf S, Lisgo SN, Vortmeyer A, Weinberger DR, Mane S, Hyde TM, Huttner A, Reimers M, Kleinman JE, Sestan N* (2011). Spatio-temporal transcriptome of the human brain. Nature. 478, 483-9. (*Co-first author)**

### 受试者须知

方案名称：人脑皮层结构的多组学研究

方案编号：KY2022-590

方案版本号：03，2022 年 7 月 29 日

知情同意书版本号：02，2022 年 7 月 29 日

研究机构：复旦大学附属华山医院神经外科、复旦大学脑科学研究院

主要研究者（负责研究医师）：陈亮

尊敬的受试者：

您将被邀请参加一项临床研究。本须知提供给您一些信息以帮助您决定是否参加此项临床研究。请您仔细阅读，如有任何疑问请向负责该项研究的研究者提出。我们将和您或您的家人进行详细沟通，向您介绍该项研究的有关情况，如果您同意参与这项研究，也请您提供与疾病有关的情况，包括发病过程、家族史、以前就诊情况及曾经做过一些检查结果等, 我们将对您进行编号，建立病历档案。

您参加本项研究是自愿的。本次研究已通过本研究机构伦理审查委员会审查。

**1. 研究概况：**

对人类和非人灵长类动物大脑的比较研究很难以传统的研究方法实现。由于伦理和法律的限制，实验生物学中使用的大多数方法不适用于研究人类和非人灵长类大脑。因此，我们对控制灵长类大脑发育和功能的分子和细胞机制的大部分理解，都来源于对一些遗传距离较远的模式生物的实验研究。虽然这些模型系统的使用使生物医学科学发生了革命性的变化，但是从它们中推断出来的结论有时候并不适用于类人猿。

近年来，测序技术的发展，为研究人脑和非人灵长类动物大脑开启了新的大门。例如通过物种间全基因组比较，发现了人类加速进化的基因和调控序列，且神经系统基因的进化突变富集于调控序列，说明人类和非人灵长类之间的许多表型差异可能来源于调控序列的变异。而调控序列的变异往往通过影响基因表达来发挥作用。因此近一步的一系列研究通过脑组织的转录组分析，寻找在人脑中特异性表达的基因及其功能。申请人前期的工作通过比较人、黑猩猩和恒河猴大脑多脑区的转录组，发现了成年人脑在分子和细胞水平上的重构，及其发育基础。

**2. 研究目的：**

本研究希望通过对成年人脑的多个脑区进行多组学测序，并与非人灵长类如猩猩、猴子进行比较，探索人脑特异性基因表达的调控机制，特别解析非编码区序列变化对基因表达的影响。

**3. 研究过程：**

本研究拟对 400 例大脑或小脑皮层有癫痫、出血、创伤或肿瘤性病灶需要切除的病人进行手术样本收集，收集后的样本用于离体电生理检测及多组学测序。您在神经外科手术中切除的组织除供临床常规病理检查之外可能还有一些剩余将丢弃，我们将收集这些剩余的组织标本约 0.5-1cm3 进行离体电生理检测及多组学测序，探索人脑特异性基因表达的调控机制。

**4. 风险与不适：**

您的手术将由专业外科医师操作，本临床研究仅仅是采集一些临床病理检查剩余的组织标本不增加任何额外风险，不影响临床治疗决策。

**5. 获益：**

通过对您的生物样本进行检测或研究可能有助于对疾病做出诊断，或为疾病的研究提供有益的信息，也有可能对您没有直接获益。

**6. 受试者职责：**

作为研究受试者，您有以下职责：提供有关自身病史和当前身体状况的真实情况；告诉研究医生自己在本次研究期间所出现的任何不适；告诉研究医生自己在最近是否曾参与其他研究，或目前正参与其他研究。

**7. 隐私问题：**

如果您决定参加本项研究，您参加研究及在研究中的个人资料均属保密。您的生物样本将以研究编号数字而非您的姓名加以标识。可以识别您身份的信息将不会透露给研究小组以外的成员，除非获得您的许可。所有的研究成员和研究申办方都被要求对您的身份保密。您的档案将保存在有锁的档案柜中，仅供研究人员查阅。为确保研究按照规定进行，必要时，政府管理部门或伦理审查委员会的成员按规定可以在研究单位查阅您的个人资料。这项研究结果发表时，将不会披露您个人的任何资料。

**8. 其他：**

您可以选择不参加本项研究，或者在任何时候通知研究者要求退出研究，您退出研究后研究者将不再收集您的数据纳入研究，您的任何医疗待遇与权益不会因此而受到影响。

如果您需要其它治疗，或者您没有遵守研究计划，或者发生了与研究相关的损伤或者有任何其它原因，研究医师可以终止您继续参与本项研究。

您可随时了解与本研究有关的信息资料和研究进展，若发生与本研究相关的安全性新信息，我们也会及时通知您。如果您有与本研究有关的问题，或您在研究过程中发生了任何不适与损伤，或有关于本项研究参加者权益方面的问题您可以通过 18221633691 与陈鸣联系。

如果您对参与本研究的权益和健康有任何问题或诉求，您可以联系本机构伦理委员会，联系电话：52888045；联系人：吴翠云。

#### 知情同意书签字页

我已经阅读了本知情同意书。

我有机会提问而且所有问题均已得到解答。

我理解参加本项研究是自愿的。

我可以选择不参加本项研究，或者在任何时候通知研究者后退出而不会遭到歧视或报复，我的任何医疗待遇与权益不会因此而受到影响。

如果我需要其它治疗，或者我没有遵守研究计划，或者发生了与研究相关的损伤或者有任何其它原因，研究医师可以终止我继续参与本项研究。

我将收到一份签过字的“知情同意书”副本。

受试者姓名：________________________

受试者签名：_________________________

日期：_______年________月________日

监护人姓名：________________________

监护人签名：_________________________

日期：______ _年_______月_______日 *（适用于受试者无认知或行为能力或为未成年人）*

见证人姓名：________________________

见证人签名：_________________________

日期：______ _年________月________日 *（适用于受试者或监护人无阅读能力，或其他需要独立见证人的情形）*

我已准确地将这份文件告知受试者，要求他/她认真阅读了这份知情同意书，对所提出的问题或疑问认真解答。

研究者姓名：________________________

研究者签名：_________________________

日期：_______年________月________日

*（注：若受试者或监护人无阅读能力，需公正的见证人签名（公正见证人，指与临床研究无关，不受临床研究相关人员不公正影响的个人，在受试者或者其监护人无阅读能力时，作为公正的见证人，阅读知情同意书和其他书面资料，并见证知情同意。）；若受试者无行为或认知能力，则需获得监护人的书面知情同意；若受试者是未成年人，应获得监护人的书面知情同意，同时应获得 8-18 岁受试者的书面知情同意和 8 岁以下受试者的口头知情同意）*
